# The MTORC1-AHR pathway sustains translation and autophagy in tumours under tryptophan stress

**DOI:** 10.1101/2023.01.16.523931

**Authors:** Pauline Holfelder, Lucas Hensen, Mirja Tamara Prentzell, Patricia Razquin Navas, Marie Solvay, Ahmed Sadik, Deepak Sayeeram, Ulrike Rehbein, Tobias Bausbacher, Anna-Sophia Egger, Leon Regin, Alexander M. Heberle, Sophie Seifert, Alexandre Leytens, Alexander Kowar, Lorenz Waltl, André Gollowitzer, Tetsushi Kataura, Seema Ouhadi, Ivana Karabogdan, Madlen Hotze, Tobias Kipura, Alienke van Pijkeren, Yang Zhang, Maria Rodriguez Peiris, Cecilia Barile, Jingjing Zhu, Lea Emmy Timpen, Florian Hatzmann, Anja Reintjes, Bianca Berdel, Luis F. Somarribas Patterson, Michèle Reil, Vera Peters, Jose Ramos Pittol, Ineke van ˈt Land-Kuper, Lara E. Eckhardt, Philipp Sievers, Felix Sahm, Shad A. Mohammed, Teresa Börding, Sönke Harder, Fabricio Loayza-Puch, Viktor I. Korolchuk, Hartmut Schlüter, Jörn Dengjel, Andreas Koeberle, Carsten Hopf, Saskia Trump, Marcel Kwiatkowski, Christine Sers, Benoit J. Van den Eynde, Christiane A. Opitz, Kathrin Thedieck

## Abstract

Tumours face tryptophan (Trp) depletion, but the mechanisms sustaining protein biosynthesis under Trp stress remain unclear. We report that Trp stress increases the levels of the translation repressor EIF4EBP1. Yet, at the same time, EIF4EBP1 is selectively phosphorylated by the metabolic master regulator MTORC1 kinase, preventing EIF4EBP1 from inhibiting translation. MTORC1 activity under Trp stress is unexpected because the absence of amino acids is typically linked with MTORC1 inhibition. EIF4EBP1-sensitive translation in Trp starved cells is sustained by EGFR and RAS signalling to MTORC1. Via this mechanism, Trp stress enhances the synthesis and activity of the aryl hydrocarbon receptor (AHR). This is noteworthy as Trp catabolites are known to activate AHR, and therefore Trp stress was previously considered to inhibit AHR. Trp stress-induced AHR enhances the expression of key regulators of autophagy, which sustains intracellular Trp levels and Trp-charged tRNAs for translation. Hence, Trp stress switches MTORC1 from its established inhibitory function into an enhancer of autophagy, acting through AHR. The clinical potential of this fundamental mechanism is highlighted by the activity of the mTORC1-AHR pathway and an autophagy signature in 20% of glioblastoma patients, opening up new avenues for cancer therapy.

## Main

Protein biosynthesis is essential for tumour survival and progression^1,2^ and requires an adequate supply of amino acids. Cancers contain poorly vascularized areas and inefficient tumour blood vessels compromise nutrient delivery. As the least abundant essential amino acid, tryptophan (Trp) will be the first to become limiting upon nutrient restriction. Trp catabolism is often upregulated in cancer including glioblastoma (GB), and activation of the aryl hydrocarbon receptor (AHR) by Trp catabolites promotes tumour progression^3–6^. In GB models, Trp levels decline with increasing distance from blood vessels^7^, and GB patients exhibit decreased Trp levels in blood and tumour tissue^8–10^. However, the mechanisms via which tumours sustain protein biosynthesis under Trp stress are poorly understood.

Translation in cancer is tightly coupled to the presence of amino acids^11^. Amino acids and growth factors activate the mechanistic target of rapamycin (MTOR) complex 1 (MTORC1) kinase^12^, which enhances translation initiation via phosphorylation of several substrates. These include an activating phosphorylation of S6 kinase (RPS6KB1) at T389^13,14^ and inhibitory phosphorylation of the translation repressive 4E binding protein (EIF4EBP) at multiple sites^15,16^. Phosphorylated EIF4EBP loses binding to the translation initiation factor 4E (EIF4E), thus enhancing EIF4E association with the translation initiation factor 4G (EIF4G)^17–19^. MTORC1 repression by amino acid limitation is well documented for arginine, leucine, methionine, glutamine and asparagine^20–23^. Relatively little is known about how Trp stress signals to MTORC1 and translation. Trp deprivation inhibits phosphorylation of RPS6KB1 at T389^24–26^, which is in line with the idea that MTORC1 activity is low and does not enhance translation. However, we find that translation under Trp stress is enabled by (1) MTORC1-mediated EIF4EBP1 phosphorylation which sustains translation initiation and (2) MTORC1-AHR-driven autophagy providing Trp for tryptophanyl-tRNA charging.

### MTORC1 phosphorylates EIF4EBP1 and sustains translation under Trp stress

Human GB tissues exhibit extensive regions of Trp restriction (**Fig. 1a**), and we wondered whether and how this aggressive tumour sustains protein biosynthesis when Trp is scarce. We compared physiological Trp levels of 78 µM^27^ to 24 h of Trp starvation in LN-18 GB cells. Trp stress reduced the amount of charged tryptophanyl-tRNA (**Fig. 1b**), suggesting an impact on translation. We conducted a puromycin incorporation assay (**Fig. 1c,d**) to assess *de novo* protein biosynthesis^28^. In line with reduced translation, Trp stress decreased puromycin incorporation, but it was even further diminished by the translation elongation inhibitor cycloheximide (CHX)^29^. Thus, protein biosynthesis under Trp stress continues at a lower level. We analysed the concentration-dependent effects of exogenous Trp and found that decreasing Trp levels reduced phosphorylation of RPS6KB1-T389 (**Fig. 1e,f**) as reported previously^24,25^, and enhanced the level of the translation repressor EIF4EBP1 (**Fig. 1e,g**). These findings are in line with overall reduced translation. We asked which mechanisms drive protein synthesis when Trp is scarce, and we made the following observations: (1) As the total levels of EIF4EBP1 increased with declining Trp concentrations, also EIF4EBP1 phosphorylation at T37/46 was induced (**Fig. 1e,h)**. Normalization of EIF4EBP1 phosphorylation to the EIF4EBP1 total levels showed that as the total levels increased upon Trp stress, phosphorylation of EIF4EBP1 increased such that the ratio between the two was maintained (**Extended Data Fig. 1a & throughout the manuscript**). We corroborated the result in LN-229 GB cells (**Extended Data Fig. 1b-e**), and we reasoned that EIF4EBP1 phosphorylation may sustain translation initiation under Trp stress. (2) As the extracellular Trp concentration dropped from 78 to 7.8 µM, the intracellular Trp concentration dropped by two orders of magnitude from 267.9 µM to 3.5 µM (**Extended Data Fig. 1f-h**). Yet, intracellular Trp declined only marginally with further extracellular Trp reduction (**Extended Data Fig. 1f-h**). This suggested that the cells sustained the low intracellular Trp concentration to secure Trp as a building block for translation.

**Figure 1:**
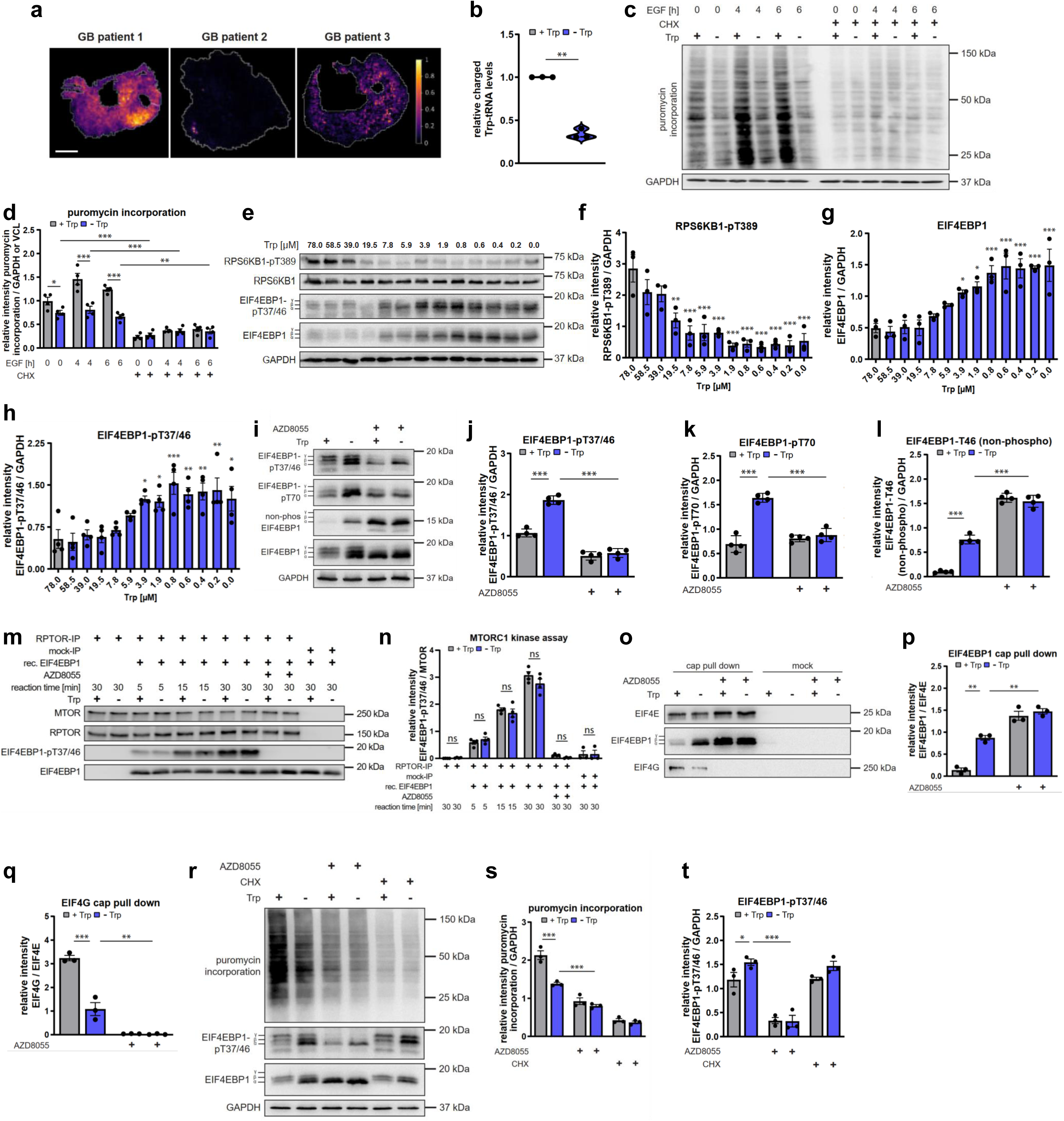
MTORC1 phosphorylates EIF4EBP1 and sustains translation under Trp stress. (a) MALDI mass spectrometry imaging (MALDI-MSI) of Trp distribution in human glioblastoma (GB) sections. Colour scale: purple, low Trp; yellow, high Trp; scale bar: 1 mm. (n = 3). (b) Tryptophanyl-tRNA charging under Trp replete conditions or Trp stress. Relative aminoacylation levels were determined by qRT-PCR using tRNA-specific primers. LN-18 cells. (n = 3). (c) Translation under Trp replete conditions or Trp stress. Puromycin (5 µg/mL, 5 min) incorporation in GB cells, unstimulated or stimulated with epidermal growth factor (EGF, 10 ng/mL, stimulation period as indicated), and treated with the translation elongation inhibitor cycloheximide (CHX) (2 µg/mL, 6.5 h). LN-18 cells. (n = 4). (d) Quantification of puromycin incorporation in (c). (e) Trp stress differentially alters signalling towards translation initiation. Trp concentration row: Cells were cultured in medium with the indicated Trp concentrations for 24 h. LN-18 cells. (n = 3-4). (f) Quantification of RPS6KB1-pT389 (S6K-pT389) in (e). (n = 3). (g) Quantification of EIF4EBP1 (4E-BP1) in (e). (n = 3). (h) Quantification of EIF4EBP1-pT37/46 (4E-BP1-pT37/46) in (e). (n = 4). (i) The MTOR inhibitor AZD8055 (100 nM, 24 h) blocks EIF4EBP1-pT37/46 (4E-BP1-pT37/46) and EIF4EBP1-pT70 (4E-BP1-pT70) and increases unphosphorylated (non-phospho) EIF4EBP1-T46 (4E-BP1-T46) under Trp stress. LN-18 cells. (n = 4). (j) Quantification of EIF4EBP1-pT37/46 (4E-BP1-pT37/46) in (i). (k) Quantification of EIF4EBP1-pT70 (4E-BP1-pT70) in (i). (l) Quantification of (non-phospho) EIF4EBP1-T46 (4E-BP1-T46) in (i). (m) Kinase assay with MTORC1 purified from LN-18 cells under Trp replete conditions or Trp stress. MTOR actively phosphorylates EIF4EBP1-pT37/46 (4E-BP1-pT37/46) upon Trp stress. Substrate: recombinant (rec.) EIF4EBP1 (4E-BP1) (100 ng). Negative controls: No EIF4EBP1 (4E-BP1), mock-IP, MTOR inhibitor AZD8055 (100 nM). Reaction time as indicated. (n = 4). (n) Quantification of EIF4EBP1-pT37/46 (4E-BP1-pT37/46) normalized to MTOR in (m). (o) Cap pull down with m7-GTP beads from LN-18 cells under Trp replete conditions or Trp stress. The MTOR inhibitor AZD8055 (100 nM, 1 h) enhances EIF4EBP1 (4E-BP1)-EIF4E binding and decreases EIF4G-EIF4E binding under Trp stress. (n = 3). (p) Quantification of EIF4EBP1 (4E-BP1) binding to EIF4E in (o). (q) Quantification of EIF4G binding to EIF4E in (o). (r) The MTOR-inhibitor AZD8055 (100 nM, 4 h) inhibits translation upon Trp stress. Puromycin assay. Translation elongation inhibitor cycloheximide (CHX) (2 µg/mL, 4 h), puromycin (5 µg/mL, 5 min). LN-18 cells. (n = 3). (s) Quantification of puromycin incorporation in (r). (t) Quantification of EIF4EBP1-pT37/46 (4E-BP1-pT37/46) in (r). Cells were cultured in the presence of Trp (+Trp, grey, 78 µM) or absence of Trp (-Trp, blue, 0 µM) for 24 h, if not indicated otherwise. One-way ANOVA followed by a Šídák’s multiple comparisons test was applied (d, f, g, h, j, k, l, n, p, q, s, t). For (b) a two-tailed paired Student’s t test was performed. Data are presented as mean ± SEM. *p < 0.05, **p < 0.01, ***p < 0.001. n.s., not significant.

We investigated how EIF4EBP1 phosphorylation is regulated and whether it is required for translation under Trp stress. Apart from MTORC1, several other kinases can phosphorylate EIF4EBP1-T37/46^30^. We inhibited MTORC1 by the ATP-analogue inhibitor AZD8055^31^, which efficiently blocks EIF4EBP1 phosphorylation by MTORC1^15,16,19^. AZD8055 reduced Trp stress-induced EIF4EBP1 phosphorylation at the MTORC1 substrate sites T37/46 and T70 (**Fig. 1i-k, Extended Data Fig. 1i,j**). In further support of MTORC1-mediated phosphorylation of EIF4EBP1 under Trp stress, an antibody recognizing non-phosphorylated EIF4EBP1-T46 detected an increased signal upon AZD8055 treatment of Trp-depleted cells (**Fig. 1i,l, Extended Data Fig. 1k**). An *in vitro* kinase assay confirmed that under Trp stress MTORC1 remained active and phosphorylated EIF4EBP1 (**Fig. 1m,n**). We conclude that MTORC1 phosphorylates EIF4EBP1 under Trp stress. EIF4EBP1 phosphorylation inhibits EIF4EBP1 binding to the translation initiation factor EIF4E, thereby enabling EIF4E-EIF4G complex formation and cap-dependent translation initiation^32^. In line with the puromycin assay (**Fig. 1c,d**), a cap binding assay (**Fig. 1o-q, Extended Data Fig. 1l-q**) confirmed that under Trp stress, translation was reduced but remained active: Trp stress enhanced EIF4EBP1-EIF4E binding (**Fig. 1o,p**) while EIF4E-EIF4G association (**Fig. 1o,q**) was preserved at a lower level. Under Trp stress, AZD8055 further enhanced EIF4EBP1-EIF4E binding (**Fig. 1o,p**) and abolished EIF4E-EIF4G complex formation (**Fig. 1o,q**), demonstrating that EIF4EBP1 phosphorylation by MTORC1 is required for cap-dependent translation under Trp stress. The small compound 4EGI-1 is an EIF4EBP1 agonist that enhances EIF4EBP1-EIF4E binding and suppresses EIF4E-EIF4G association, thus inhibiting translation initiation at the cap^33,34^. Under Trp stress, both 4EGI-1 (**Extended Data Fig. 1r,s**) and AZD8055 (**Fig. 1r-t, Extended Data Fig. 1t**) inhibited puromycin incorporation, further supporting that EIF4EBP1 phosphorylation by MTORC1 sustains translation under Trp stress. Our data show that MTORC1-mediated phosphorylation prevents the Trp stress-induced increase in total EIF4EBP1 from inhibiting translation, thereby sustaining protein biosynthesis. We conclude that MTORC1 is active under Trp stress, which expands the common view that MTORC1 is inhibited by amino acid deprivation^35–38^ and puts Trp into a unique position in the control of MTORC1.

### The EGF receptor and RAS signal Trp stress to EIF4EBP1

Growth factors activate MTORC1 via class I phosphoinositide 3-kinases (PI3Ks)^36^. The pan class I PI3K inhibitor Pictilisib (GDC-0941)^39^ inhibited EIF4EBP1 phosphorylation at T37/46 in Trp-restricted cells (**Extended Data Fig. 2a-c**), indicating that PI3K signals to MTORC1 and EIF4EBP1 when Trp is scarce. We went on to investigate which upstream cues mediate Trp stress signalling to EIF4EBP1. PI3K is a key effector of the small GTPase RAS^40^, whose activation has been primarily assigned to growth factor inputs^41,42^. RAS activation by stress is less established^43–46^, and nutrient stress or Trp restriction have so far not been linked to RAS. In a RAS-GTP pull down assay^44^, Trp restriction enhanced RAS binding to a RAF-RAS-binding domain (GST-RAF1), indicative of enhanced RAS-GTP loading and activity (**Fig. 2a,b**). Knockdown of all RAS isoforms (KRAS/HRAS/NRAS) reduced phosphorylation of EIF4EBP1-T37/46 in Trp-deprived cells (**Fig. 2c-e, Extended Data Fig. 2d**). The epidermal growth factor (EGF) receptor (EGFR) acts upstream of RAS^47^ and is frequently amplified in GB^48^. Autophosphorylation of the EGFR at Y1068^49,50^ was enhanced in Trp-deprived cells with (**Extended Data Fig. 2e,f)** and without EGF stimulation (**Fig. 2f,g)**. Trp stress enhanced EGFR internalization to perinuclear endosomes (**Fig. 2h,i)**, consistent with EGFR activation^51^. The pan-ERBB (EGF receptor family) inhibitor Afatinib^52^ as well as the EGFR-specific inhibitor Erlotinib^53^ reduced Trp stress-induced phosphorylation of EIF4EBP1-T37/46 (**Fig. 2j-l, Extended Data Fig. 2g)**, showing that EGFR mediates Trp stress signalling to EIF4EBP1. EGFR activation by Trp stress without exogenous EGF addition (**Fig. 2f,g)** suggested a contribution by an endogenous ligand. Whereas *EGF* mRNA levels were reduced by Trp restriction (**Fig. 2m**), levels of *EREG* (epiregulin) mRNA (**Fig. 2n**) as well as unglycosylated and glycosylated pro-EREG proteins^54^ were enhanced with declining Trp levels (**Fig. 2o-q**). We conclude that EGFR and RAS drive signalling to EIF4EBP1 in Trp-deprived cells. Given the activation of the EGFR-RAS pathway, one would have anticipated both *bona fide* MTORC1 substrates EIF4EBP1 and RPS6KB1 to become phosphorylated. Surprisingly, however, Trp stress exerted opposing effects on the two MTORC1 substrates as it enhanced phosphorylation of EIF4EBP1, but reduced phosphorylation of RPS6KB1 (**Fig. 1e,f,h, Extended Data Fig. 1b,c,e**). We found that this divergent regulation was mediated by the MTORC1 suppressor Sestrin2 (SESN2)^55–60^. Trp stress induced SESN2 levels, and SESN2 knockdown selectively enhanced RPS6KB1-T389 phosphorylation (**Fig. 2r-u, Extended Data Fig. 2h**). Thus, SESN2 represses RPS6KB1 phosphorylation but not EIF4EBP1 phosphorylation under Trp stress.

**Figure 2:**
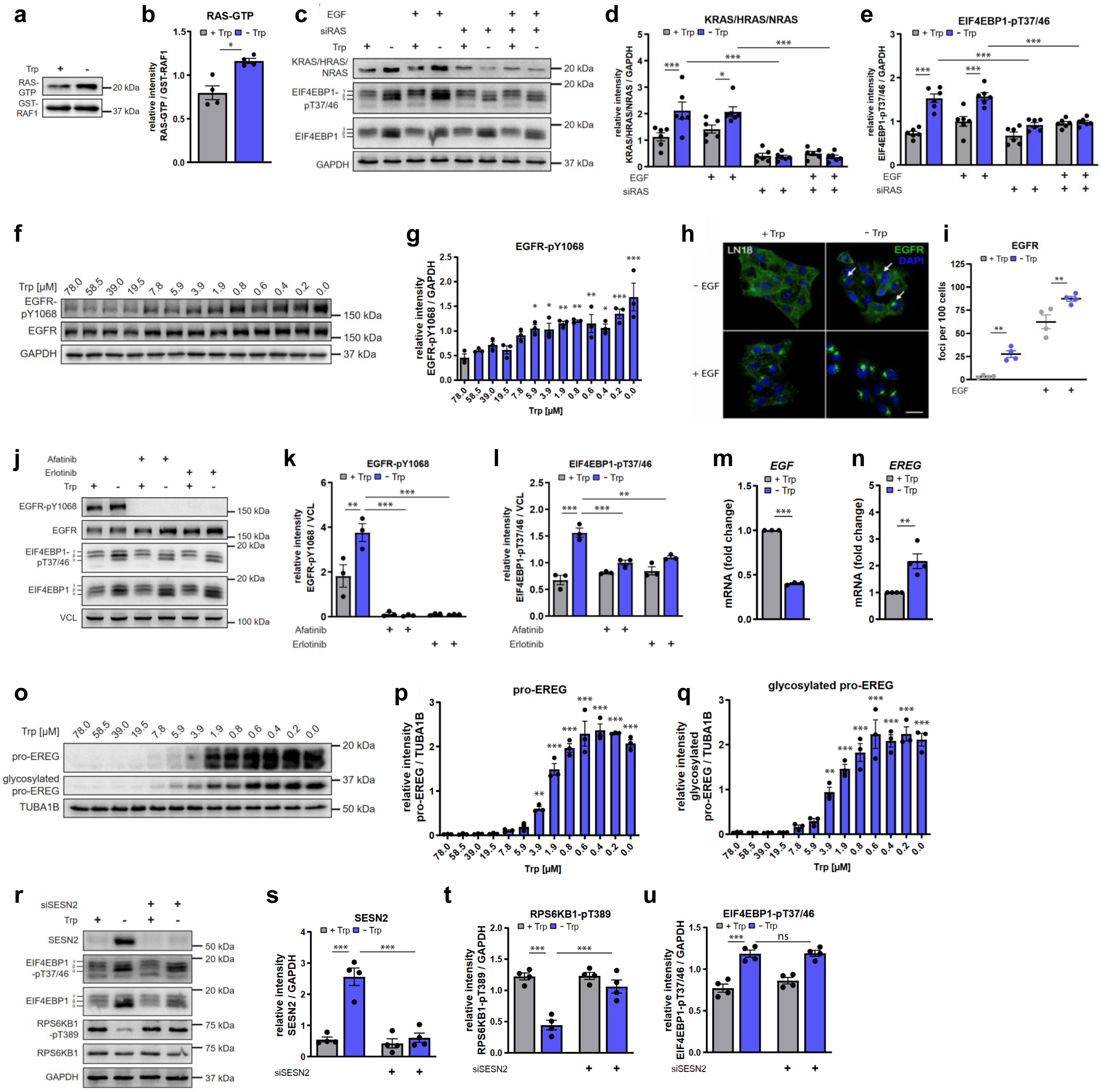
The EGF receptor and RAS signal Trp stress to EIF4EBP1. (a) Trp stress enhances RAS-GTP binding to RAF1-GST. RAS activity was measured using GST-coupled RAF-RAS–binding domain pull down experiments. Cells treated with EGF (10 ng/mL, 30 min). LN-18 cells. (n = 4). (b) Quantification of RAS-GTP in (a). (c) Pan-RAS (KRAS/HRAS/NRAS) knockdown (siRAS) reduces EIF4EBP1-pT37/46 (4E-BP1-pT37/46) induction by Trp stress. Cells were either unstimulated or stimulated with EGF (10 ng/mL, 15 min). LN-18 cells. (n = 6). (d) Quantification of KRAS/HRAS/NRAS in (c). (e) Quantification of EIF4EBP1-pT37/46 (4E-BP1-pT37/46) in (c). (f) Trp deprivation gradually enhances EGFR autophosphorylation, similar to the induction of EIF4EBP1-pT37/46 (4E-BP1-pT37/46). Trp concentration row: Cells were cultured in medium with the indicated Trp concentrations for 24 h. Detections of the same samples as in Figure 1e. LN-18 cells. (n = 3). (g) Quantification of EGFR-pY1068 in (f). (h) Trp stress enhances EGFR internalization to perinuclear endosomes. Immunofluorescence of EGFR localization. Cells were unstimulated or stimulated with EGF (10 ng/mL, 15 min). EGFR, green; nucleus, blue (DAPI). Scale bar: 10 µm. LN-18 cells. (n = 4). (i) Quantification of EGFR foci per 100 cells in (h). (j) The pan-ERBB receptor inhibitor Afatinib (10 µM, 1 h) and the EGFR-specific inhibitor Erlotinib (10 µM, 1 h) both inhibit Trp restriction-induced EIF4EBP1-pT37/46 (4E-BP1-pT37/46). Cells were stimulated with EGF (10 ng/mL, 30 min). LN-18 cells. (n = 3). (k) Quantification of EGFR-pY1068 in (j). (l) Quantification of EIF4EBP1-pT37/46 (4E-BP1-pT37/46) in (j). (m) Trp stress reduces transcripts of the EGFR ligand EGF. *EGF* mRNA relative to *18S rRNA* measured by qRT-PCR. LN-18 cells. (n = 3). (n) Trp stress enhances transcripts of the EGFR ligand Epiregulin (EREG). *EREG* mRNA relative to *18S rRNA* measured by qRT-PCR. LN-18 cells. (n = 4). (o) Trp deprivation gradually enhances EREG expression, similar to the induction of EGFR autophosphorylation and EIF4EBP1-pT37/46 (4E-BP1-pT37/46). Trp concentration row: Cells were cultured in medium with the indicated Trp concentrations for 24 h. LN-18 cells. (n = 3). Detections of the same samples as in Figure 1e. (p) Quantification of pro-EREG in (o). (q) Quantification of glycosylated pro-EREG in (o). (r) SESN2 knockdown rescues repressed RPS6KB1-pT389 (S6K-pT389) levels upon Trp stress, but does not affect EIF4EBP1-pT37/46 (4E-BP1-p37/46). LN-229 cells. (n = 4). (s) Quantification of SESN2 in (r). (t) Quantification of RPS6KB1-pT389 (S6K-pT389) in (r). (u) Quantification of EIF4EBP1-pT37/46 (4E-BP1-pT37/46) in (r). Cells were cultured in the presence of Trp (+Trp, grey, 78 µM) or absence of Trp (-Trp, blue, 0 µM) for 24 h, if not indicated otherwise. One-way ANOVA followed by a Šídák’s multiple comparisons test was applied (d, e, g, i, k, l, p, q, s, t, u). For (b,m,n) a two-tailed paired Student’s t test was performed. Data are presented as mean ± SEM. *p < 0.05, **p < 0.01, ***p < 0.001. n.s., not significant.

### EIF4EBP1-sensitive translation induces AHR expression and activity under Trp stress

We explored the protein repertoire, which is induced by Trp stress. 364 proteins were increased upon Trp deprivation (**Fig. 3a**). Ribosome profiling showed ribosome pausing at Trp codons (TGG) under Trp stress (**Extended Data Fig. 3a**), potentially leading to accumulation of incomplete polypeptides. However, the proteome data showed that peptide coverage was not altered by Trp stress (**Extended Data Fig. 3b,c**) and extended beyond Trp residues (**Extended Data Fig. 3d**), supporting that Trp stress did not interrupt translation prematurely. We compared the proteomes under Trp stress, generalized amino acid stress in amino acid-free DMEM or HBSS media, or starvation of methionine, another essential amino acid required for translation initiation (**Fig. 3b,c,d**). Trp stress shared only 29 upregulated proteins with general amino acid starvation (**Extended Data Fig. 3e**), and 31 proteins with methionine stress (**Extended Data Fig. 3f**). 335 or 333 upregulated proteins were exclusive to Trp stress, as compared to generalized amino acid or methionine stress, respectively (**Extended Data Fig. 3e,f**). Similar results were found for the downregulated proteins in either condition (**Extended Data Fig. 3g,h**). Neither general amino acid stress nor methionine stress enhanced EIF4EBP1 phosphorylation (**Fig. 3e-i, Extended Data Fig. 3i**). Thus, Trp stress has a profound impact on the proteome, which differs from generalized amino acid stress and methionine deprivation.

**Figure 3:**
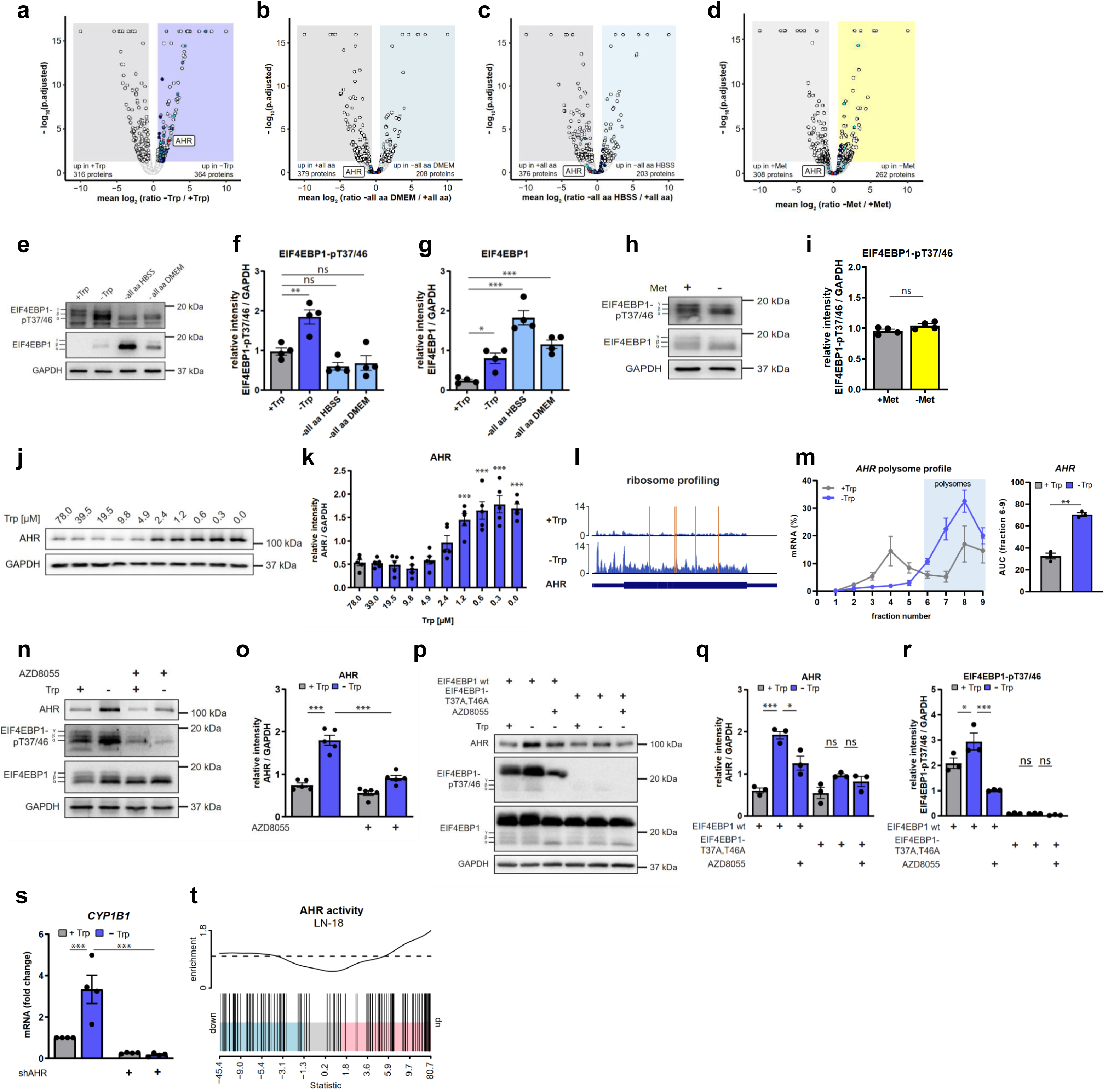
EIF4EBP1-sensitive translation induces AHR expression and activity under Trp stress. (a) The Trp stress proteome reveals an increase of the aryl hydrocarbon receptor (AHR). Volcano plot of relative protein abundances. The Trp replete condition (+Trp) was compared to Trp stress (-Trp) for 24 h. The AHR is marked in red, the other colours correspond to the GO terms in Fig. 4a. LN-18 cells. (n = 3). Full proteome and list of marked proteins are in Table S1. (b) The proteome upon generalized amino acid stress in DMEM medium shows no increase of the AHR. Volcano plot of relative protein abundances. The amino acid replete condition (+all aa) was compared to withdrawal of all amino acids in DMEM medium (-all aa DMEM) for 24 h. The AHR is marked in red, the other colours correspond to the GO terms in Fig. 4a. LN-18 cells. (n = 4). Full proteome and list of marked proteins are in Table S1. (c) The proteome upon generalized amino acid stress in HBSS medium shows no increase of the AHR. Volcano plot of relative protein abundances. The amino acid replete condition (+all aa) was compared to withdrawal of all amino acids in HBSS (-all aa HBSS) for 24 h. The AHR is marked in red, the other colours correspond to the GO terms in Fig. 4a. LN-18 cells. (n = 4). Full proteome and list of marked proteins are in Table S1. (d) The methionine (Met) stress proteome shows no increase of the AHR. Volcano plot of relative protein abundances. The methionine replete condition (+Met) was compared to Met stress (-Met). The AHR is marked in red, the other colours correspond to the GO terms in Fig. 4a. LN-18 cells. (n = 4). Full proteome and list of marked proteins are in Table S1. (e) Hyperphosphorylation of EIF4EBP1 (4E-BP1) at T37/46 is specific to Trp stress. Cells were cultured under amino acid replete conditions in DMEM (control, +Trp), under Trp stress (-Trp), or under withdrawal of all amino acids in HBSS (-all aa HBSS) or DMEM (-all aa DMEM) for 24 h. LN-18 cells. (n = 4). (f) Quantification of EIF4EBP1-pT37/46 (4E-BP1-pT37/46) in (e). (g) Quantification of EIF4EBP1 (4E-BP1) in (e). (h) Hyperphosphorylation of EIF4EBP1 (4E-BP1) is specific to Trp stress. Cells were cultured under methionine replete conditions (control, +Met) or methionine stress (-Met) for 24 h. LN-18 cells. (n = 4). (i) Quantification of EIF4EBP1-pT37/46 (4E-BP1-pT37/46) in (h). (j) Trp deprivation gradually enhances AHR expression. Trp concentration row: Cells were cultured in medium with the indicated Trp concentrations for 24 h. LN-229 cells. (n = 5). (k) Quantification of AHR in (j). (l) AHR translation is enhanced upon Trp stress. Ribosome profiling: Ribosome protected fragment (RPF) read density is shown on the AHR transcript in the presence and absence of Trp. Reads per transcript normalized to total number of reads are shown on the y-axis. Bottom panel, short rectangles represent untranslated regions, tall rectangle indicates coding sequence. LN-229 cells. (n = 3). Replicates are shown in Extended Data Figure 3j. (m) AHR translation is enhanced upon Trp stress. Polysome profiling: Polysome profiles of AHR upon presence and absence of Trp. Analysis of relative *AHR* mRNA levels per fraction via qRT-PCR. Fraction numbers 1-5 indicate low-molecular weight polysomes, fraction numbers 6-9 indicate high-molecular weight polysomes (actively translating fractions, highlighted in light blue). LN-229 cells. (n = 3). On the right: Quantification of the area under the curve (AUC) of *AHR* in fractions 6-9. (n) The MTOR inhibitor AZD8055 (100 nM, 24 h) suppresses AHR induction by Trp stress. LN-229 cells. (n = 5). (o) Quantification of AHR in (n). (p) The MTOR inhibitor AZD8055 (100 nM, 24 h) fails to inhibit AHR levels in cells overexpressing a non-phosphorylatable EIF4EBP1 T37/46A mutant. LN-229 cells. (n = 3). (q) Quantification of AHR in (p). (r) Quantification of EIF4EBP1-pT37/46 (4E-BP1-pT37/46) in (p). (s) Short hairpin-mediated knockdown of the AHR ablates the induction of the AHR target gene *CYP1B1* by Trp stress. mRNA expression normalized to *18S rRNA*. qRT-PCR. LN-18 cells. (n = 4). (t) RNAseq analysis reveals an enhanced transcriptional AHR activity signature upon Trp stress. Barcode plots showing the status of AHR activity in cells starved of Trp for 24 h. LN-18 cells. (n = 4). The x-axis represents the values of moderated t-statistic values for all genes in the comparison. The blue and pink coloured segments represent the lower and upper quartiles of all the genes. The vertical barcode lines represent the distribution of the genes. The worm line above the barcode shows the relative density of the AHR-signature genes, which represents the direction of regulation. Cells were cultured in the presence of Trp (+Trp, grey, 78 µM) or absence of Trp (-Trp, blue, 0 µM) for 24 h, if not indicated otherwise. One-way ANOVA followed by a Šídák’s multiple comparisons test was applied (f, g, k, o, q, r, s). For (i) and (m) a two-tailed paired Student’s t test was performed. Data are presented as mean ± SEM. *p < 0.05, **p < 0.01, ***p < 0.001. n.s., not significant.

Intriguingly, the AHR was specifically induced in the Trp stress proteome (**Fig. 3a)** and AHR levels increased with declining Trp concentrations (**Fig. 3j,k**). We investigated if AHR induction was mediated by EIF4EBP1-sensitive translation. Ribosome profiling showed an increased association of ribosomes with AHR transcripts under Trp stress as well as full ribosome coverage up to the 3’ end (**Fig. 3l, Extended Data Fig. 3j**). Also, polysome profiling indicated that the AHR was preferentially translated under Trp stress (**Fig. 3m**). In agreement, the translation elongation inhibitor CHX fully suppressed AHR protein induction by Trp stress (**Extended Data Fig. 3k,l**) and AHR induction upon Trp stress was EGFR-(**Extended Data Fig. 3m-p**), MTOR-(**Fig. 3n,o, Extended Data Fig. 3q,r**) and EIF4EBP1-sensitive **(Extended Data Fig. 3s,t)**. Trp stress enhanced AHR levels only in cells expressing EIF4EBP1 wildtype but not in cells expressing a non-phosphorylatable EIF4EBP1-T37/46A mutant^15^ (**Fig. 3p-r, Extended Data Fig. 3u**). Furthermore, AZD8055 could not reduce AHR levels in cells with non-phosphorylatable EIF4EBP1-T37/46A, showing that MTORC1 controls AHR expression via EIF4EBP1.

Notably, Trp stress enhanced not only AHR levels but also AHR activity, as determined by induction of the AHR target gene *CYP1B1* that was suppressed by AHR knockdown (**Fig. 3s, Extended Data Fig. 3v**). Also, RNAseq analysis (**Table S2**) revealed that Trp stress enhanced a transcriptional AHR activity signature^61^ (**Fig. 3t, Extended Data Fig. 3w-y**). This finding was unexpected as the AHR is typically considered to be activated by Trp metabolites^62–65^, but not under Trp restriction when Trp metabolites are low or absent.

### The MTORC1-AHR pathway enhances autophagy to replenish intracellular Trp levels and to sustain tryptophanyl-tRNA charging under Trp stress

We went on to investigate the functions of the MTORC1-AHR axis under Trp stress. Trp limitation enhanced the enrichment of gene ontology (GO) terms related to macropinocytosis and lysosomes (**Fig. 4a, Table S1**). Macropinocytosis is a non-selective process driven by the EGFR and RAS^66,67^, leading to uptake of the extracellular fluid-phase and macromolecules^68^. We measured internalization of fluorescently-labeled dextran to stain macropinosomes and found them increased upon Trp depletion (**Fig. 4b,c**). Upon macropinosome maturation, their content is delivered to the lysosomal compartment for degradation^68^. In agreement with the GO term enrichment of lysosomal components in the Trp stress proteome (**Fig. 4a**), the LAMP2 (lysosomal associated membrane protein 2)-positive lysosomal area (**Fig. 4d,e**) and lysosome activity (**Fig. 4f,g**) were enhanced by Trp stress. Autophagy constitutes a lysosome function, which is critical in conjunction with macropinocytosis to break down macromolecules and fuel cancer metabolism under nutrient limitation^68–70^. Lipidated LC3 (ubiquitin-like protein microtubule associated protein 1 light chain 3 beta, MAP1LC3B), termed LC3-II, decorates autophagosomes and serves as an anchor for autophagy receptor proteins^71^. To assess autophagic flux^72^, LC3-II was detected in conjunction with inhibition of lysosome acidification by Bafilomycin A_1_ (BafA) (**Fig. 4h-k, Extended Data Fig. 4a**). Under Trp replete conditions, BafA only mildly increased LC3-II, indicative of low autophagic flux, and AHR inhibition by stemreginin-1 (SR1)^73^ did not affect LC3-II levels. MTOR inhibition enhanced LC3-II, consistent with its well-known function as an autophagy suppressor under nutrient replete conditions^72^. Trp stress enhanced autophagic flux, based on LC3-II induction by BafA (**Fig. 4h,i**) and an increased autophagy mRNA signature^74^ (**Fig. 4l, Extended Data Fig. 4b-c**). Under Trp stress, MTOR inhibition as well as AHR inhibition reversed the LC3-II level back to that without BafA (**Fig. 4h,i**). Also, the number of foci of the autophagy receptor SQSTM1 (sequestosome 1, p62)^72^, an alternative readout for autophagy, was reduced by MTOR inhibition as well as by AHR inhibition under Trp stress (**Fig. 4m,n**). We conclude that the MTORC1-AHR pathway drives autophagy under Trp stress. This finding is intriguing as MTORC1 switches from its canonical, inhibitory role in autophagy to becoming an autophagy activator under Trp stress.

**Figure 4:**
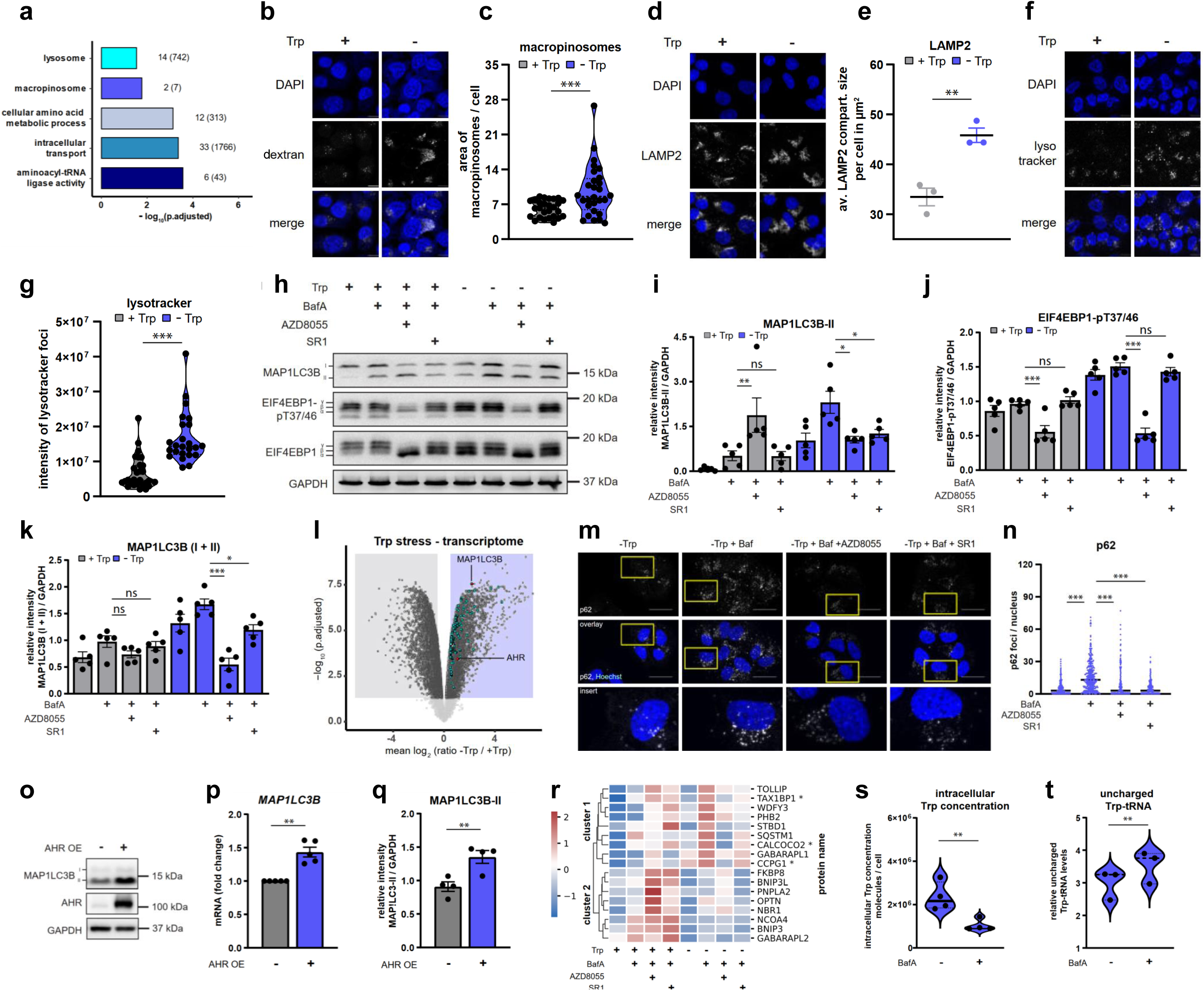
The MTORC1-AHR pathway enhances autophagy to replenish intracellular Trp levels and to sustain tryptophanyl-tRNA charging under Trp stress. (a) Gene Ontology (GO) terms related to macropinocytosis and lysosomes are enriched in the Trp stress proteome. GO term enrichment analysis of proteins upregulated under Trp stress (-Trp) in Figure 3a. Proteins were considered to be upregulated with FC ≥ 1.5 and an adjusted p-value < 0.05. The length of the bar represents the log10 Benjamini-Hochberg corrected p-value. Indicated for each term is the number of associated proteins in the Trp stress proteome; in brackets: total number of associated proteins per term. Proteins that belong to the GO terms are marked in the corresponding colours in the volcano plots in Figure 3a-d and are listed in Table S1. (b) Trp stress increases the area of intracellular macropinosomes. Uptake assay of fluorescently labelled 70 kDa - dextran (dextran, white) in cells stained with DAPI (blue), stimulated with EGF (10 ng/mL, 30 min). Scale bar: 10 µm. LN-18 cells. (n = 4). (c) Quantification of the area of macropinosomes per cell in (b). (d) Trp stress increases lysosomal compartment size. Immunofluorescence staining of LAMP2 (white) in cells stained with DAPI (blue), stimulated with EGF (10 ng/mL, 15 min). Scale bar: 10 µm. LN-18 cells. (n = 4). (e) Quantification of the area of LAMP2 staining (lysosomal compartment size) per cell in µm^2^ (d). (f) Trp stress increases lysosomal activity. Live cell imaging of LysoTracker Red DND-99 (lysotracker, 30 min, white) in cells stained with DAPI (blue). Scale bar: 10 µm. LN-18 cells. (n = 5). (g) Quantification of the intensity of lysotracker foci in (f). (h) Trp stress enhances autophagic flux (BafA-induced MAP1LC3B lipidation), which is suppressed by inhibition of MTOR or the AHR. BafA (100 nM, 2 h), MTOR inhibitor AZD8055 (100 nM, 24 h), AHR inhibitor SR1 (1 µM, 24 h). LN-18 cells. (n = 5). (i) Quantification of lipidated MAP1LC3B-II (LC3-II) in (h). (j) Quantification of EIF4EBP1-pT37/46 (4E-BP1-pT37/46) in (h). (k) Quantification of MAP1LC3B total levels (LC3 I + II) in (h). (l) Volcano plot showing differentially regulated genes in the Trp stress (-Trp) versus the Trp replete condition (+Trp) in RNAseq data of LN-18 cells. Autophagy-related genes are coloured in cyan. AHR and MAP1LC3B (LC3) are shown in red (n = 4). Analysis of the same dataset as in Figure 3t. Table S2 lists all the differentially expressed autophagy-related genes. (m) MTOR or AHR inhibition reduces autophagy upon Trp stress. Immunofluorescence staining of p62 foci. BafA (100 nM, 2 h), MTOR inhibitor AZD8055 (100 nM, 24 h), AHR inhibitor SR1 (1 µM, 24 h). Scale bar: 20 µm. LN-18 cells. (n = 4). (n) Quantification of p62 foci in (m). (o) AHR overexpression (OE) increases AHR and MAP1LC3B (LC3) protein. LN-18 cells. (n = 4). (p) AHR overexpression (OE) increases *MAP1LC3B (LC3)* mRNA levels. mRNA expression normalized to *18S rRNA*. qRT-PCR. LN-18 cells. (n = 5). (q) Quantification of MAP1LC3B (LC3) in (o). (r) Heatmap of autophagy regulators measured by targeted quantitative proteomics in cells cultured in the presence or absence of Trp, treated without or with BafA (100 nM, 2 h), and with BafA in combination with the MTOR inhibitor AZD8055 (100 nM, 24 h) or the AHR inhibitor SR1 (1 µM, 24 h). LN-18 cells. Colours represent z-scored averages of the relative abundances n > 4 replicates. (n = 5, except for +Trp, +BafA, +SR1 condition with n = 4). Peptides were normalized to their respective heavy-labelled spiked-in peptides. Proteins labelled with an asterisk show a significant difference (p-value < 0.05) between the -Trp BafA condition and the -Trp BafA AZD8055 condition. (s) Cells under Trp stress exhibit a further decrease in intracellular Trp when lysosomal function is inhibited. Intracellular Trp concentrations (molecules Trp / cell) in cells treated with and without BafA (100 nM, 2 h). LN-18 cells. (n = 4). The -Trp condition already shown in Extended Data Fig. 1g. (t) BafA (100 nM, 2 h) increases the levels of uncharged tryptophanyl-tRNAs upon Trp stress. Relative aminoacylation levels were determined by qRT–PCR using tRNA-specific primers. LN-18 cells. (n = 3). The -Trp condition is also shown in Fig. 1b. Cells were cultured in the presence of Trp (+Trp, grey, 78 µM) or absence of Trp (-Trp, blue, 0 µM) for 24 h, if not indicated otherwise. One-way ANOVA followed by a Šídák’s multiple comparisons test was applied (i, j, k). For (c, e, g, p, q, s, t) a two-tailed paired Student’s t test was performed. Data are presented as mean ± SEM. *p < 0.05, **p < 0.01, ***p < 0.001. n.s., not significant.

Under nutrient replete conditions, MTORC1 inhibits autophagy by repressing the kinase ULK1 (unc-51 like autophagy activating kinase 1) and the TFEB/ TFE3 (transcription factor EB/ transcription factor binding to IGHM enhancer 3) transcription factors, two central enhancers of autophagy^72^. These mechanisms cannot provide an explanation for the observed reduction in autophagy by MTOR inhibition under Trp stress: if the MTOR inhibitor were to further decrease ULK1 and TFEB/TFE3 phosphorylation, it would enhance autophagy, contrary to our observation. We therefore reasoned that under Trp stress, the MTORC1-AHR pathway must induce autophagy via another route. Of note, Trp stress enhanced not only LC3 lipidation but also LC3 total levels in an MTOR-and AHR-dependent manner (**Fig. 4k**). In the Trp stress transcriptome, LC3 was among the most upregulated transcripts (**Fig. 4l, Table S2**). In agreement with AHR-driven LC3 expression, we identified an AHR binding motif in the LC3 promotor region (**Extended Data Fig. 4d**). Ribosome profiling showed that also LC3 translation was enhanced under Trp stress (**Extended Data Fig. 4e**). Thus, LC3 is transcribed and translated *de novo* under Trp stress. AHR overexpression (**Fig. 4o-q, Extended Data Fig. 4f,g**) was sufficient to enhance *LC3* at mRNA levels (**Fig. 4p**) and LC3 lipidation (**Fig. 4o,q**) demonstrating that the transcription factor AHR induces LC3 expression and autophagy. To investigate a wider array of autophagy regulators under Trp stress (**Fig. 4l**), we quantified a panel of core autophagy regulators by targeted quantitative proteomics upon lysosome blockade by BafA in combination with MTOR or AHR inhibition (**Fig. 4r**). Under Trp replete conditions, most proteins were enhanced by MTOR inhibition, in line with increased autophagic flux. In contrast, under Trp stress several central autophagy receptor, anchor and cargo proteins including GABARAPL1 (GABA type A receptor associated protein like 1) and SQSTM1 (p62) depended on MTOR and AHR (**Fig. 4r**, cluster 1). Taken together, the MTORC1-AHR pathway induces the expression of core components of the autophagy machinery and drives autophagic flux under Trp stress. Via this mechanism, Trp stress triggers a functional switch of MTORC1 in autophagy, making it an autophagy activator. Of note, BafA-mediated inhibition of lysosomal function induced a drop in intracellular Trp (**Fig. 4s**) and increased the proportion of uncharged tryptophanyl-tRNAs (**Fig. 4t**) in Trp-deprived cells. This indicates that lysosome-derived Trp helps cancer cells to overcome Trp limitation by maintaining tryptophanyl-tRNA charging for translation.

### The mTORC1-AHR pathway is active in human glioblastomas

We next interrogated whether the MTORC1-AHR pathway is present in human tumours. We clustered transcriptome data of human GB (The Cancer Genome Atlas, TCGA)^75^ based on the transcripts regulated by Trp stress in the GB cell lines (**Table S2**), yielding seven patient subgroups. Using an AHR activity signature^61^ we identified two patient subgroups (1 and 2) with high AHR activity scores (**Fig. 5a**). Patient subgroup 1 (∼10% of the GB patients) exhibited low AHR mRNA levels, whereas patient subgroup 2 (∼20% of the GB patients) showed high *AHR* mRNA levels (**Fig. 5b**). We reasoned that the latter patient subgroup may have experienced Trp restriction, as Trp stress induces AHR expression. The TCGA reverse phase protein array (RPPA) data revealed that subgroup 2, with high AHR expression, exhibited higher EIF4EBP1-T37/46 phosphorylation than subgroup 1 **(Fig. 5c**), whereas RPS6KB1-T389 phosphorylation was similar in both patient groups **(Fig. 5d**). This further supports that GB tumour tissues in subgroup 2 show signs of Trp deprivation, as they featured low RPS6KB1 and high EIF4EBP1 phosphorylation as well as high AHR levels. In agreement, subgroup 2 showed enhanced expression of a transcriptional autophagy regulator signature^74^ (**Fig. 5e**). Thus, an augmented autophagy signature associates with high EIF4EBP1 phosphorylation, high AHR levels, and high AHR activity. Taken together, the data suggest that the MTORC1-AHR pathway is active and drives autophagy in 20% of human GB. The AHR positively regulates various enzymes of the ceramide biosynthetic pathway, including sphingomyelin phosphodiesterases (SMPD, sphingomyelinases)^76,77^ that catalyze the conversion of sphingomyelin to ceramides in lysosomes^78^. Therefore, we investigated ceramides as potential markers for the activity of the MTORC1-AHR pathway, detectable in tumour tissue. In GB cells, Trp stress increased the levels of ceramides, which were suppressed by MTOR and AHR inhibitors (**Fig. 5f, Table S3**). Thus, MTOR and the AHR enhance ceramide levels upon Trp stress, further highlighting the MTORC1-AHR pathway as a driver of lysosomal function. In GB tissues, MALDI-MS imaging revealed that regions of high Trp and high ceramides were mutually exclusive (**Fig. 5g**), suggesting that low Trp and high ceramides – marking the start and endpoints of the MTORC1-AHR pathway – are coupled in human tumours and indicate active MTORC1-AHR signalling.

**Figure 5:**
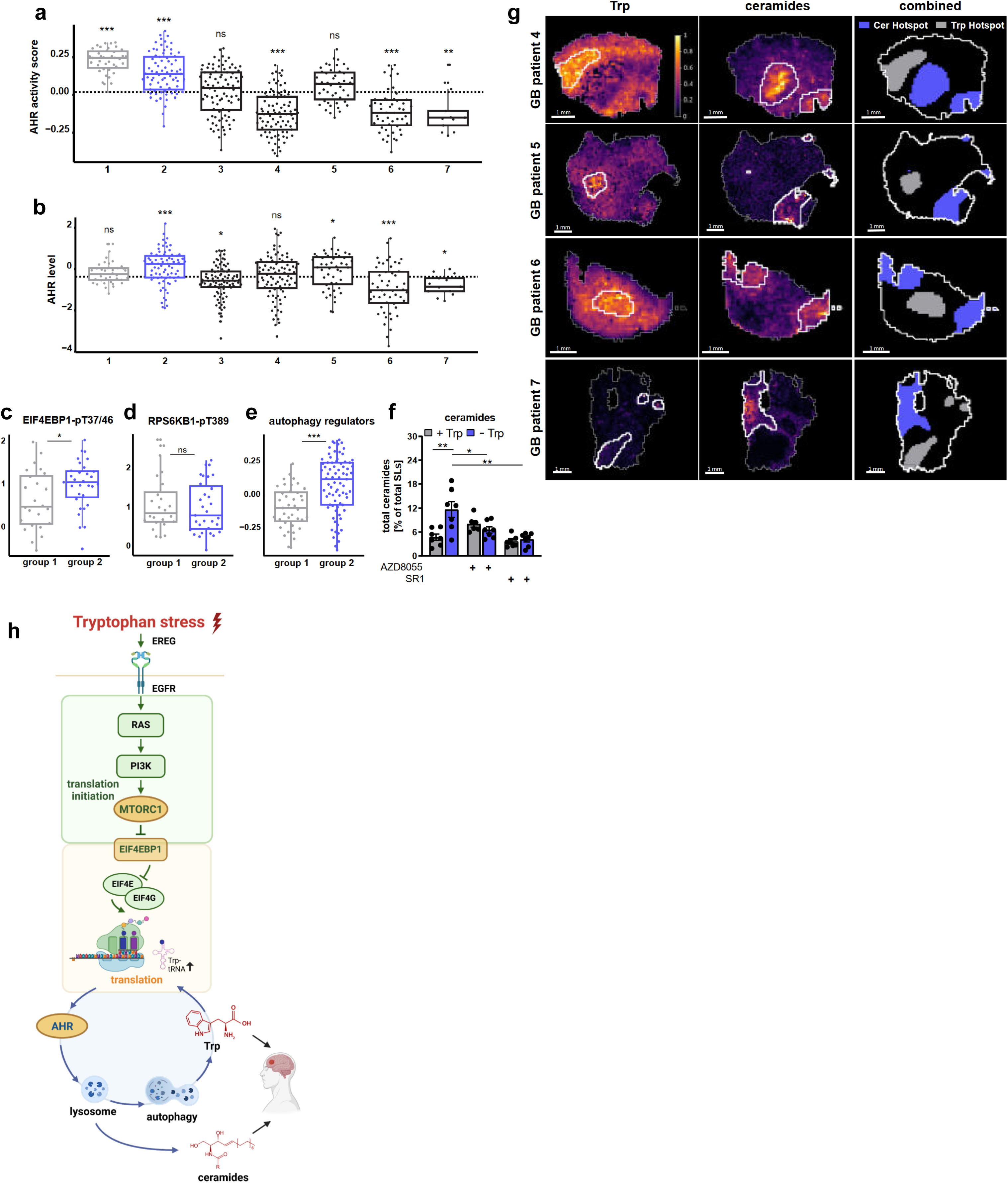
The mTORC1-AHR pathway is active in human glioblastomas. (a, b) Boxplot representation of the distribution of the (a) AHR activity score and (b) the normalized expression values of AHR in the seven glioblastoma patient subgroups of human GB (The Cancer Genome Atlas, TCGA). The black dotted line represents the mean AHR activity score or normalized AHR expression values across all patient samples. The p-values were determined based on comparing the average score or expression to the corresponding mean of all patient sample groups. (c-e) Boxplot representation of the distribution of TCGA RPPA data of EIF4EBP1-pT37/46 (4E-BP1-pT37/46) (c), P70S6K1-pT389 (S6K-pT389) (d), and the enrichment score of autophagy regulators (e), between groups 1 (high AHR activity and low AHR mRNA expression; grey) and 2 (high AHR activity and high AHR mRNA expression; blue). (f) Trp stress enhances the cellular proportion of ceramides in an MTOR- and AHR-dependent manner. Ratio of ceramides to total sphingolipids (SLs). MTOR inhibitor AZD8055 (100 nM, 24 h), AHR inhibitor SR1 (1 µM, 24 h). LN-18 cells. (n = 7). (g) In human GB, regions of high Trp and high ceramides are mutually exclusive. MALDI-MSI of Trp and ceramide hotspots and their intersections in human GB samples. (n = 4). Scale bar: 1 mm. (h) Schematic representation of the MTORC1-AHR pathway. Cells were cultured in the presence of Trp (+Trp, grey, 78 µM) or absence of Trp (-Trp, blue, 0 µM) for 24 h. A two-tailed unpaired Student’s t test was performed in (f). For bioinformatic analysis, statistic is described in the Method Details section (a-e). Data are presented as mean ± SEM. *p < 0.05, **p < 0.01, ***p < 0.001. n.s., not significant.

## Discussion

We address the fundamental question of how tumours maintain protein biosynthesis when Trp is scarce (**Fig. 5h**). We report that under Trp stress EIF4EBP1 phosphorylation by MTORC1 sustains translation initiation, and EIF4EBP1-controlled translation induces the AHR. As a result, MTORC1-AHR signalling emerges as a novel pathway under Trp stress, which drives autophagy, replenishing intracellular Trp levels and tryptophanyl-tRNAs for translation. Hence, the MTORC1-AHR pathway enables protein synthesis under Trp stress in a twofold manner, at the level of translation initiation and by providing tryptophanyl-tRNAs. The MTORC1-AHR pathway features several molecular functions that differ from the established picture of MTORC1 or the AHR alone. Trp stress enhances the level of the key translation repressor EIF4EBP1. At the same time, however, MTORC1 remains active and selectively phosphorylates EIF4EBP1, which expands the common view that amino acid limitation inhibits MTORC1^79,80^. EIF4EBP1-sensitive translation promotes expression of the AHR, known to be activated by Trp via its catabolites^62–65^. However, we find that also Trp stress activates the AHR. The MTORC1-AHR pathway induces major controllers of autophagy. Therefore, Trp stress switches MTORC1’s role in autophagy from its canonical inhibitory function to becoming an autophagy enhancer, whereby MTORC1 acts via the AHR.

Our finding that MTORC1 remains active when the essential amino acid Trp is depleted changes our view on the interplay of amino acids with this key tumour driver. Withdrawal of the single amino acids arginine, leucine, methionine, glutamine, and asparagine represses MTORC1-dependent translation through inhibition of the RRAG GTPases^20,79^ and other lysosomal regulators^21–23^. Trp stress is different in that it selectively enhances phosphorylation of EIF4EBP1, but not RPS6KB1, and the Trp stress proteome profoundly differs from other amino acid stresses. Selective inhibition of RPS6KB1 by the RRAG repressor SESN2^55,56^ has not been described so far^79,80^, and it requires further investigation whether the lower affinity of RPS6KB1 to MTORC1^13,19,81^ explains RPS6KB1’s higher sensitivity to MTORC1 inhibition by SESN2. Long term leucine or arginine starvation has been linked with MTORC1 activation via PI3K and AKT^82^, however there was no divergence between RPS6KB1 and EIF4EBP1 phosphorylation, and autophagy was down, suggesting that induction of the MTORC1-AHR pathway is Trp-stress-specific. MTORC1 has also been found activated by long term glutamine deprivation^83,84^ but this is mediated by amino acid transporters enhancing the influx of amino acids^83^, which differs from Trp stress induction of the EGFR-RAS axis upstream of MTORC1. Given that EGFR and RAS also are upstream of the MAPK pathway, which impinges on EIF4E-driven translation in tumours^1,2,30^, MAPKs may contribute to the MTORC1-mediated Trp stress response upstream of translation.

Like MTORC1, translation initiation is generally considered as being inhibited by nutrient starvation and stress^85^. We report that under Trp stress, translation is reduced but remains active. Thus, inhibitory and activating cues balance translation under Trp restriction, allowing for expression of a Trp stress-specific protein repertoire. Trp stress-induced proteome remodelling may have evolved as Trp is the physiologically least abundant amino acid, and a drop in Trp levels is an early indicator of an upcoming starvation for all amino acids. In other words, the Trp stress-sensitive MTORC1-AHR pathway likely serves as a sentinel mechanism that senses an imminent decline in amino acids. By adapting its translation repertoire, the cell can express proteins that are necessary to cope with nutrient starvation while most amino acids are still sufficiently available. The integrated stress response mediated via the GCN2-EIF2A-ATF4 pathway restricts translation initiation under Trp stress^30,86,87^ and it will be intriguing to address its interplay with MTORC1-EIF4EBP1 mediated translation. Incorporation of phenylalanine (Phe) instead of Trp has recently been suggested to sustain translation in Trp-restricted tumours^88^. There was a low overall frequency of such events in our proteome data, and only two peptides harbouring a Trp-Phe exchange increased upon Trp stress (**Table S1**). Thus, mobilization of Trp by autophagy appears sufficient to sustain translation of the Trp-containing proteome, including the AHR.

Clinical trials with AHR inhibitors are currently ongoing for cancer immunotherapy^89^. Our findings demonstrate a novel role for AHR in tumour cells, i.e. enabling them to cope with Trp stress by enhanced autophagy. Like the TFEB/TFE transcription factors, which enhance lysosome biogenesis and autophagy^72^, also the AHR enhances autophagy by mediating the expression of a differential autophagy signature that encompasses key components of the autophagy machinery including LC3, GABARAPL1, and SQSTM1 (p62). However, unlike TFEB/TFE3, AHR is activated by MTORC1. This is how MTORC1 switches its role and actively contributes to enhanced autophagy under Trp stress.

Trp restriction has been suggested earlier to inhibit the AHR in tumours because Trp metabolites enhance AHR activity^90^. However, our data suggest that this is not a good strategy as Trp stress enhances AHR levels and activity. Rather, reduced Trp levels and enhanced levels of Trp catabolites synergize in boosting AHR activity and its oncogenic outcomes. Indeed, Trp depletion potentiates the effect of Trp catabolites as AHR ligands^91^ and promotes Treg differentiation^92^. Our finding that Trp stress activates the AHR highlights the need to not only consider Trp metabolites but also Trp levels to predict AHR activity in tumours. Our data suggest that the MTORC1-AHR pathway is active in 20% of GB patients, attesting to the clinical potential of this fundamental mechanism. These patients may benefit from autophagy suppression by inhibitors of MTORC1 and its upstream cues as well as its novel downstream target AHR^93,94^. As Trp shortage occurs in many cancers^6,63^, the MTORC1-AHR pathway may also be active in tumour entities beyond GB. Tumour metabolites attract growing attention as predictive markers, and we determine low Trp and high ceramide levels to be at the start and end points of the MTORC1-AHR pathway. The Trp/ceramide ratio hence warrants clinical testing as a predictive marker for response to drugs targeting the MTORC1-AHR pathway.

---

**Extended Data Figure 1:**
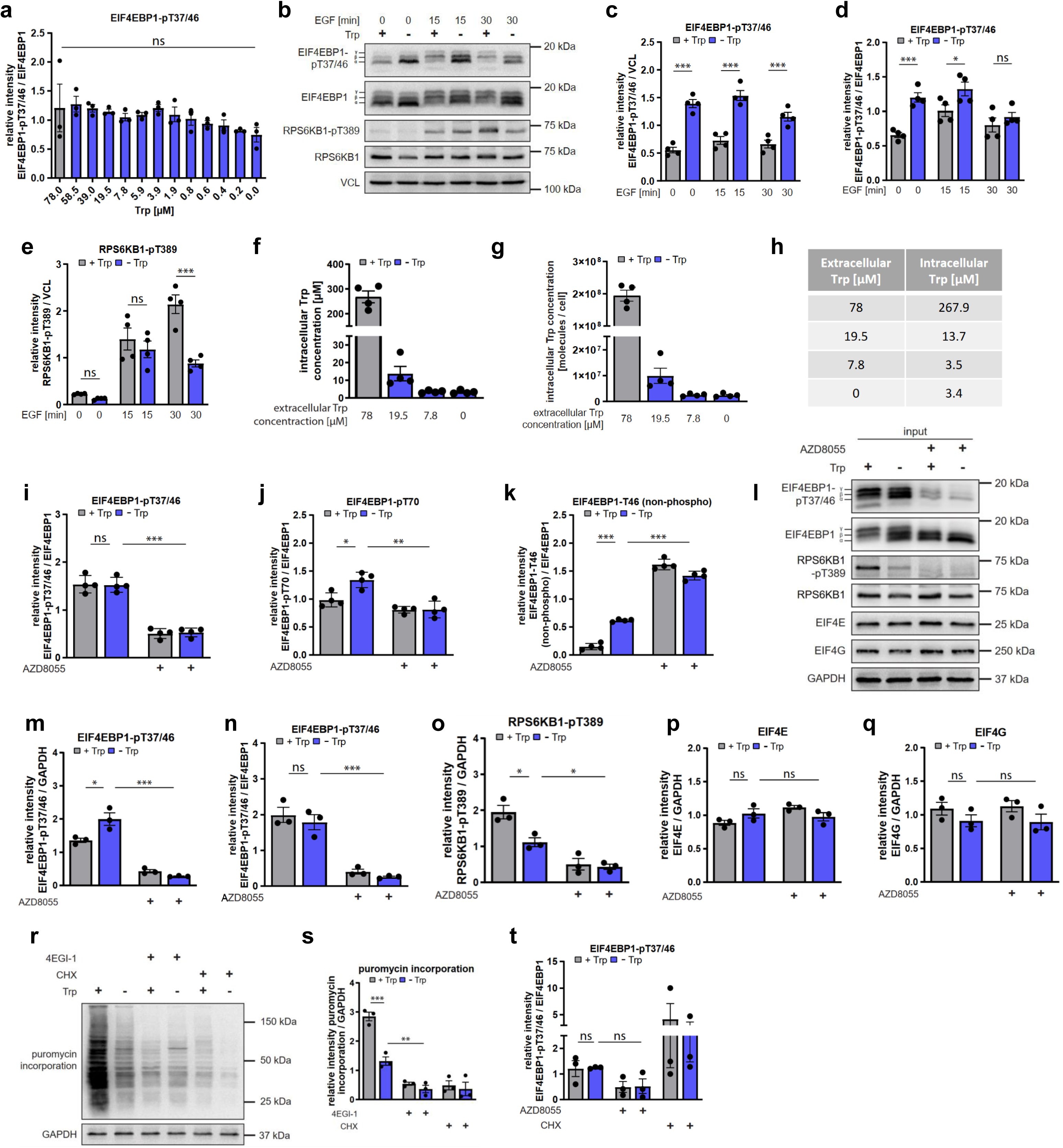
Under Trp stress the intracellular Trp concentration is sustained. (a) Quantification of EIF4EBP1-pT37/46 (4E-BP1-pT37/46) over total EIF4EBP1 (4E-BP1) in Figure 1e. (b) Trp stress differentially alters signalling towards translation initiation also in LN-229 cells. Cells were stimulated with EGF (10 ng/mL) for the indicated time periods. (n = 4). (c) Quantification of EIF4EBP1-pT37/46 (4E-BP1-pT37/46) in (b). (d) Quantification of EIF4EBP1-pT37/46 (4E-BP1-pT37/46) over total EIF4EBP1 (4E-BP1) in (b). (e) Quantification of RPS6KB1-pT389 (S6K-pT389) in (b). (f) Intracellular Trp concentrations in µM. Cells were cultured in medium with the indicated Trp concentrations for 24 h. LN-18 cells. (n = 4). (g) Intracellular Trp concentrations in molecules per cell. Cells were cultured in medium with the indicated Trp concentrations for 24 h. LN-18 cells. (n = 4). (h) Comparison between extracellular and intracellular Trp concentrations in µM. Cells were cultured in medium containing 78, 19.5, 7.8 or 0 µM Trp for 24 h. Cultivation with 7.8 or 0 µM Trp reduces the intracellular Trp concentration from 267.9 µM to 3.5-3.4 µM. LN-18 cells. (n = 4). (i) Quantification of EIF4EBP1-pT37/46 (4E-BP1-pT37/46) over total EIF4EBP1 (4E-BP1) in Figure 1i. (j) Quantification of EIF4EBP1-pT70 (4E-BP1-pT70) over total EIF4EBP1 (4E-BP1) in Figure 1i. (k) Quantification of EIF4EBP1-pT46 (non-phospho) (4E-BP1-pT46) over total EIF4EBP1 (4E-BP1) in Figure 1i. (l) Input for cap pull down in Figure 1o. (m) Quantification of EIF4EBP1-pT37/46 (4E-BP1-pT37/46) in (l). (n) Quantification of EIF4EBP1-pT37/46 (4E-BP1-pT37/46) over total EIF4EBP1 (4E-BP1) in (l). (o) Quantification of RPS6KB1-pT389 (S6K-pT389) in (l). (p) Quantification of EIF4E in (l). (q) Quantification of EIF4G in (l). (r) The EIF4EBP1 (4E-BP1) agonist 4EGI-1 (10 µM, 24 h) inhibits translation upon Trp stress. Puromycin assay. Translation elongation inhibitor cycloheximide (CHX) (2 µg/mL, 24 h), puromycin (5 µg/mL, 5 min). LN-18 cells. (n = 3). (s) Quantification of puromycin incorporation in (r). (t) Quantification of EIF4EBP1-pT37/46 (4E-BP1-pT37/46) over total EIF4EBP1 (4E-BP1) in Figure 1r. Cells were cultured in the presence of Trp (+Trp, grey, 78 µM) or absence of Trp (-Trp, blue, 0 µM) for 24 h, if not indicated otherwise. One-way ANOVA followed by a Šídák’s multiple comparisons test was applied in (a, i, j, k, m, n, o, p, q, s, t), two-way ANOVA followed by a Šídák’s multiple comparisons test was applied in (c, d, e). Data are presented as mean ± SEM. *p < 0.05, **p < 0.01, ***p < 0.001. n.s., not significant.

**Extended Data Figure 2:**
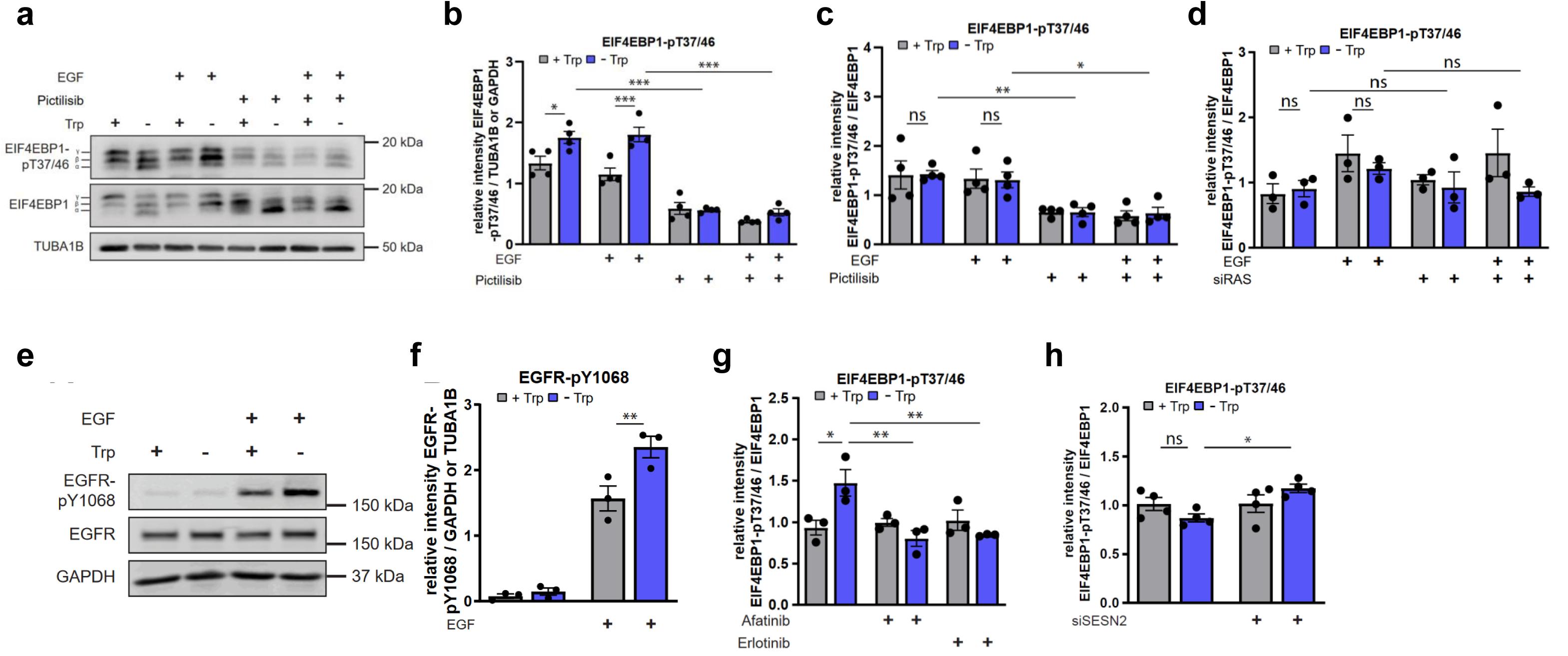
PI3K inhibition blocks Trp stress-induced EIF4EBP1 phosphorylation. (a) The class I PI3K inhibitor Pictilisib (GDC0941) (1 µM, 1 h) blocks Trp stress-induced EIF4EBP1-pT37/46 (4E-BP1-pT37/46). Cells were unstimulated or stimulated with EGF (10 ng/mL, 15 min). LN-18 cells. (n = 4). (b) Quantification of EIF4EBP1-pT37/46 (4E-BP1-pT37/46) in (a). (c) Quantification of EIF4EBP1-pT37/46 (4E-BP1-pT37/46) over EIF4EBP1 (4E-BP1) in (a). (d) Quantification of EIF4EBP1-pT37/46 (4E-BP1-pT37/46) over EIF4EBP1 (4E-BP1) in Figure 2c. (e) Trp deprivation enhances autophosphorylation of the EGFR at Y1068. Cells were unstimulated or stimulated with EGF (10 ng/mL, 15 min). LN-18 cells. (n = 3). Detections of the same samples as in (a) (lanes 1 to 4). (f) Quantification of EGFR-pY1068 in (e). (g) Quantification of EIF4EBP1-pT37/46 (4E-BP1-pT37/46) over EIF4EBP1 (4E-BP1) in Figure 2j. (h) Quantification of EIF4EBP1-pT37/46 (4E-BP1-pT37/46) over EIF4EBP1 (4E-BP1) in Figure 2r. Cells were cultured in the presence of Trp (+Trp, grey, 78 µM) or absence of Trp (-Trp, blue, 0 µM) for 24 h. One-way ANOVA followed by a Šídák’s multiple comparisons test was applied (b, c, d, f, g, h). Data are presented as mean ± SEM. *p < 0.05, **p < 0.01, ***p < 0.001. n.s., not significant.

**Extended Data Figure 3:**
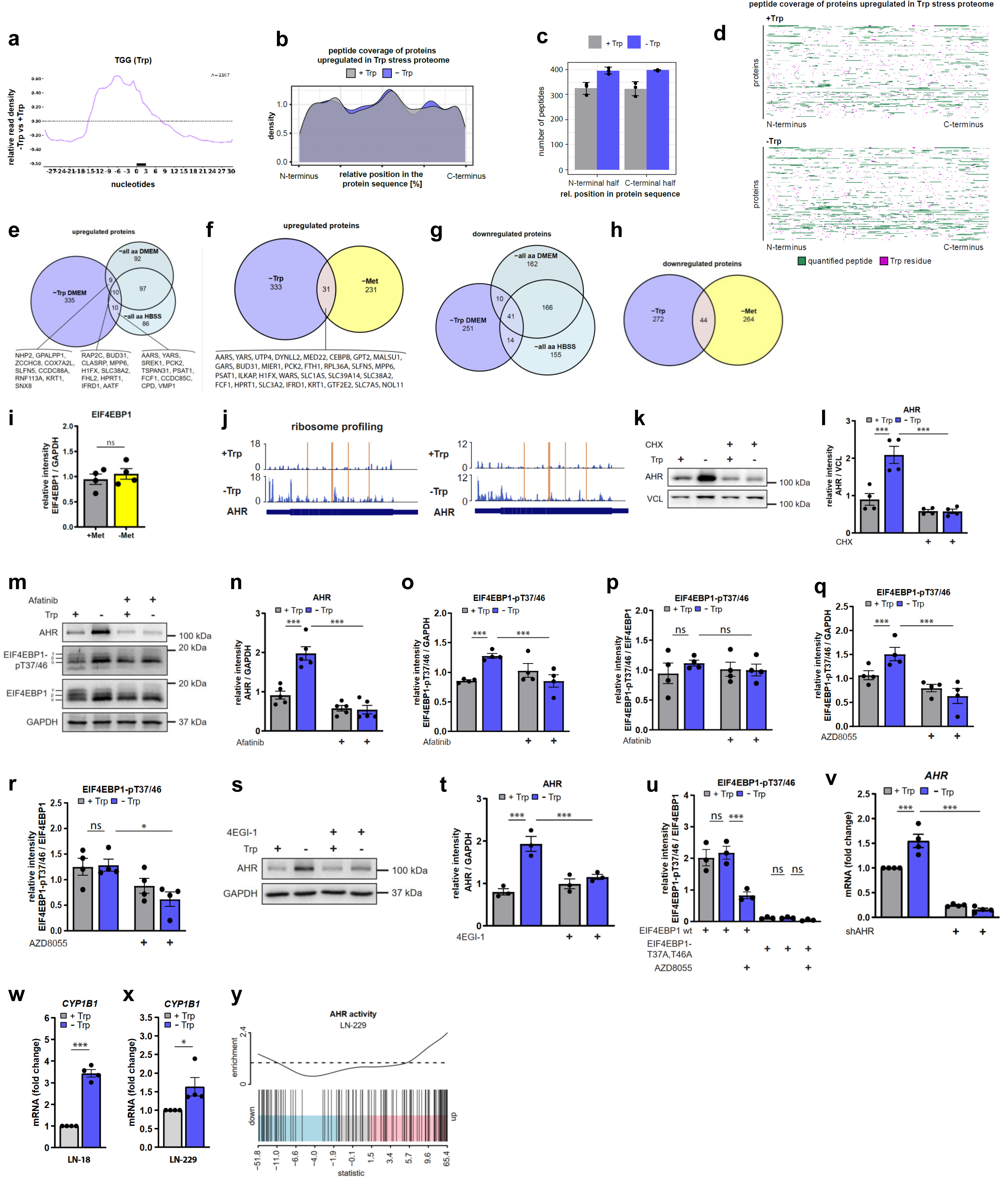
Proteins in the Trp stress proteome are fully translated. (a) Ribosome profiling: Transcriptome-wide RPF density at Trp codons (TGG) under Trp replete conditions or Trp stress. The peak indicates the accumulation of ribosomes in a window of -/+ 30 nucleotides at Trp codons under Trp stress. LN-229 cells. (n = 1). (b-d) Sequence coverage of proteins upregulated in the Trp stress proteome. (b) Density plot: relative distribution of quantified peptides from the N- to the C-terminus of proteins under Trp replete conditions (+Trp, grey) or Trp stress (-Trp, blue). No shift in the distribution of quantified peptides was observed, indicating that there was no increase in premature translation arrest events under Trp stress. (c) Distribution of quantified peptides between the N-terminal and C-terminal half. Proteins upregulated in the stress proteome were analyzed under Trp replete conditions (+Trp) and Trp stress (-Trp). (d) Peptide distribution map for all proteins under Trp replete conditions (upper panel) and Trp stress (lower panel). Relative positions of quantified peptides (green) and Trp residues (magenta) from N- to C-terminus (%) are depicted. Trp stress does not shift the peptide distribution towards the N-terminus, further suggesting that Trp stress does not result in premature translation arrest. (e) The Trp stress proteome exhibits low overlap with generalized amino acid stress in DMEM or HBSS media. Venn-diagram of upregulated proteins in Figures 3a, b and c. (f) The Trp stress proteome exhibits low overlap with methionine stress. Venn-diagram of upregulated proteins in Figures 3a and d. (g) The Trp stress proteome has low overlap with generalized amino acid stress in DMEM or HBSS media. Venn-diagram of downregulated proteins in Figures 3a, b and c. (h) The Trp stress proteome has low overlap with methionine stress. Venn-diagram of downregulated proteins in Figures 3a and d. (i) Quantification of EIF4EBP1 (4E-BP1) in Figure 3h. (j) AHR translation is enhanced upon Trp stress. Two biological replicates of the ribosome profiling data shown in Figure 3l. Ribosome protected fragment (RPF) read density is shown on the AHR transcript in the presence and absence of Trp. Reads per transcript normalized to total number of reads are shown on the y-axis. Bottom panel, short rectangles represent untranslated regions, tall rectangle indicates coding sequence. RPF data show ribosome coverage under Trp stress to be enhanced up to the 3’ end. At the Trp codons (depicted as orange lines) the read numbers increase, indicative of ribosome pausing, but this does not lead to premature translation stop. LN-229 cells. (k) Cycloheximide (CHX) (5 µg/mL, 24 h) suppresses AHR induction by Trp stress. LN-229 cells. (n = 4). (l) Quantification of AHR in (k). (m) The pan-ERBB inhibitor Afatinib (10 µM, 24 h) suppresses AHR induction by Trp stress. LN-229 cells. (n = 4-5). (n) Quantification of AHR in (m). (o) Quantification of EIF4EBP1-pT37/46 (4E-BP1-pT37/46) in (m). (p) Quantification of EIF4EBP1-pT37/46 (4E-BP1-pT37/46) over EIF4EBP1 (4E-BP1) in (m). (q) Quantification of EIF4EBP1-pT37/46 (4E-BP1-pT37/46) in Figure 3n. (r) Quantification of EIF4EBP1-pT37/46 (4E-BP1-pT37/46) over EIF4EBP1 (4E-BP1) in Figure 3n. (s) The EIF4EBP1 (4E-BP1) agonist 4EGI-1 (10 µM, 24 h) suppresses AHR induction by Trp stress. LN-229 cells. (n = 3). (t) Quantification of AHR in (s). (u) Quantification of EIF4EBP1-pT37/46 (4E-BP1-pT37/46) over EIF4EBP1 (4E-BP1) in Figure 3p. (v) Short hairpin-mediated knockdown of the AHR in Figure 3s. mRNA expression normalized to *18S rRNA*. qRT-PCR. LN-18 cells. (n = 4). (w-x) The AHR is active upon Trp stress as determined by the induction of the AHR target gene *CYP1B1*. (w) *CYP1B1* mRNA expression measured relative to *18S rRNA* by qRT-PCR, validation of RNASeq data shown in Figure 3t. LN-18 cells. (n = 4). (x) *CYP1B1* mRNA expression measured relative to *18S rRNA* by qRT-PCR, validation of RNASeq data shown in (y). LN-229 cells. (n = 4). (y) RNAseq analysis reveals an enhanced transcriptional AHR activity signature upon Trp stress. Barcode plots showing the status of AHR activity in cells starved of Trp for 24 h. LN-229 cells. (n = 4). The x-axis represents the values of moderated t-statistic values for all genes in the comparison. The blue and pink coloured segments represent the lower and upper quartiles of all the genes. The vertical barcode lines represent the distribution of the genes. The worm line above the barcode shows the relative density of the AHR-signature genes, which represents the direction of regulation. Cells were cultured in the presence of Trp (+Trp, grey, 78 µM) or absence of Trp (-Trp, blue, 0 µM) for 24 h, if not indicated otherwise. One-way ANOVA followed by a Šídák’s multiple comparisons test was applied (l, n, o, p, q, r, t, u, v), for (i, w, x) a two-tailed paired Student’s t test was performed. Data are presented as mean ± SEM. *p < 0.05, ***p < 0.001. n.s., not significant.

**Extended Data Figure 4:**
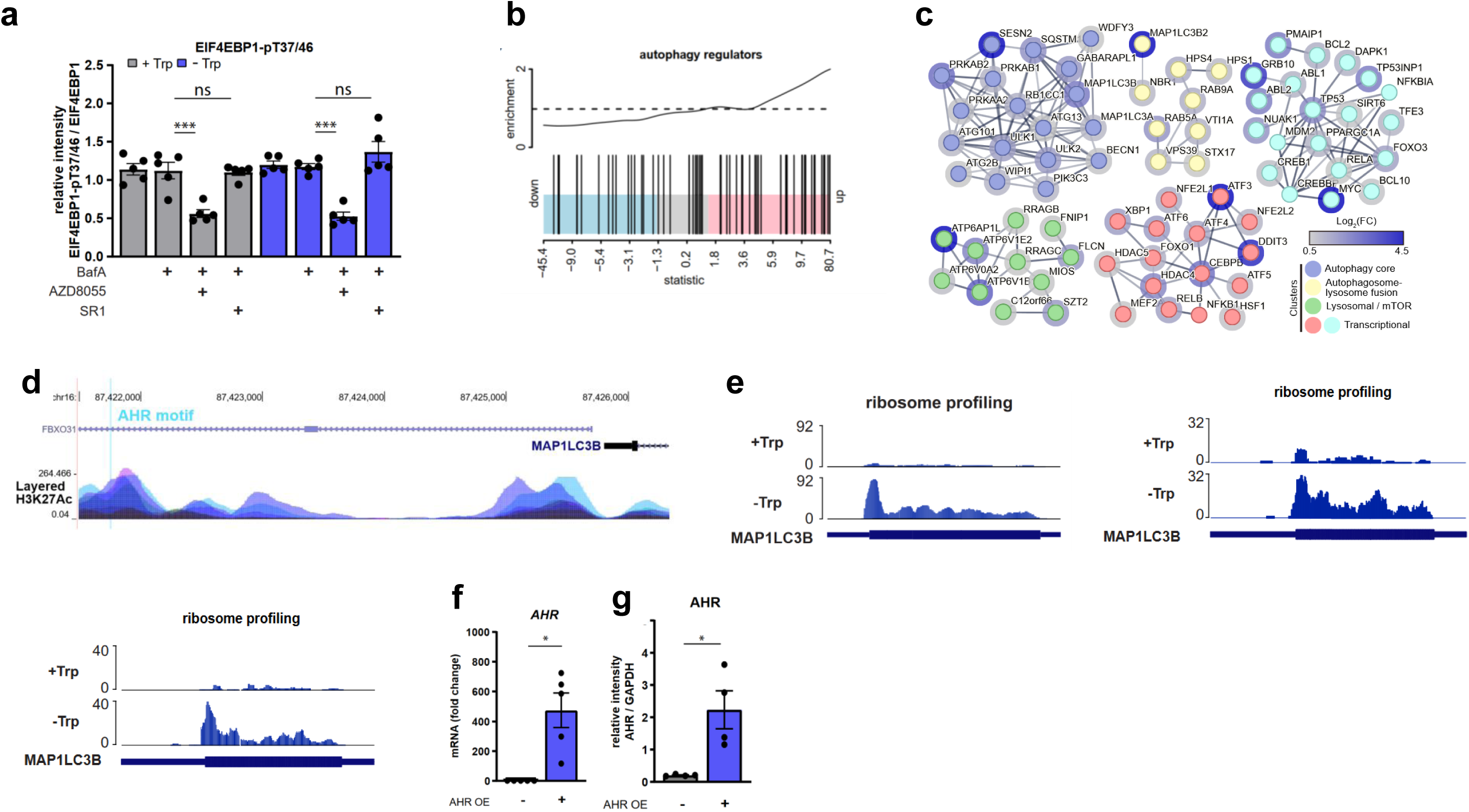
The MTORC1-AHR pathway enhances autophagy. (a) Quantification of EIF4EBP1-pT37/46 (4E-BP1-pT37/46) over total EIF4EBP1 (4E-BP1) in Figure 4h. (b) An autophagy mRNA signature is enriched upon Trp stress. Barcode plot showing enrichment of autophagy regulators in RNAseq data of cells starved of Trp for 24 h. LN-18 cells (n = 4). Analysis of the same dataset as in Figure 4l. (c) Interaction network analysis of reported physical interactions between autophagy-related proteins identified in Figure 4l. Clusters generated by k-means clustering were arbitrarily classified based on their functions. (d) AHR binding motif in the *MAP1LC3B* promoter region. The transcription factor target gene database shows an AHR binding motif (V_AHR_01_M00139, chr16: 87421754-87421771) upstream of the MAP1LC3B promoter, which is located in a region of H3K27 acetylation enrichment, indicative of active chromatin. (e) MAP1LC3B (LC3) translation is enhanced upon Trp stress. Ribosome profiling: Ribosome protected fragment (RPF) read density is shown on the MAP1LC3B (LC3) transcript in the presence and absence of Trp. Reads per transcript normalized to total number of reads are shown on the y-axis. LN-229 cells. (n = 3). Bottom panel, short rectangles represent untranslated regions, tall rectangle indicates coding sequence. RPF data show ribosome coverage under Trp stress to be enhanced up to the 3’ end. (f) AHR overexpression (OE) increases *AHR* mRNA levels. mRNA expression normalized to *18S rRNA*. qRT-PCR. LN-18 cells. (n = 5). (g) Quantification of AHR in Figure 4o. Cells were cultured in the presence of Trp (+Trp, grey, 78 µM) or absence of Trp (-Trp, blue, 0 µM) for 24 h, if not indicated otherwise. One-way ANOVA followed by a Šídák’s multiple comparisons test was applied in (a). For (f, g) a two-tailed paired Student’s t test was performed. Data are presented as mean ± SEM. *p < 0.05, ***p < 0.001. n.s., not significant.

## Methods

### Cell culture and treatments

Glioblastoma cell lines LN-18 and LN-229 were obtained from ATCC. Cells were cultured in DMEM (Biotech, P04-03600) supplemented with 10% FBS (Gibco, 10270106) and 3 mM L-glutamine (Gibco, 25030-024) or in phenol red-free DMEM (Gibco, 31053-028) supplemented with 10% FBS (Gibco, 10270106) and 2 mM L-glutamine (Gibco, 25030-024), 1 mM sodium pyruvate (Gibco, 11360039), 100 U/mL penicillin and 100 µg/mL streptomycin (Gibco, 15140122). All cell lines were maintained at 37 °C and 5% CO_2_ and regularly tested for mycoplasma contamination.

#### Cell seeding

For Ras pull down, 6.5 x 10^5^ LN-18 cells were seeded per 15 cm dish. For cap pull down, 4 x 10^6^ LN-18 cells were seeded per 15 cm dish (TPP, 93150), six dishes per condition. For polysome profiling 1.6 x 10^6^ LN-229 cells were seeded per 15 cm dish (TPP, 93150). For the ribosome profiling, 6.5 x 10^4^ LN-229 cells per mL were seeded per 15 cm dish (TPP, 93150). For the proteome analysis 3 x 10^6^ LN-18 cells were seeded per 10 cm dish (TPP, 93100). For tRNA aminoacylation assay, 3 x 10^6^ LN-18 cells were seeded per 10 cm dish (TPP, 93100). For MTOR pathway and simultaneous amino acid - sphingolipid analysis, 1 x 10^6^ LN-18 or LN-229 cells were seeded per 6 cm dish (TPP, 93060). For siRAS experiments, 2.5 x 10^5^ LN-18 cells were seeded per 6 cm dish (TPP, 93060). For analysis of AHR expression and activation, as well as RNA sequencing 4 x 10^5^ LN-229 cells per well were seeded in 6-well plates. For siRNA-mediated knockdown of SESN2 1,6 x 10^5^ cells per 6-well were seeded. For AHR overexpression 3 x 10^5^ LN-18 cells per well were seeded in 6-well plates. For EGFR immunofluorescence experiments, 1 x 10^5^ LN-18 cells were seeded per well into an 8-well IbiTreat µ-slide (Ibidi, 80826). For all other immunofluorescence experiments, 1 x 10^5^ LN-18 cells were seeded on coverslips (Hecht assistant, 41014). For the uptake assay and lysosome tracking, 5 x 10^4^ LN-18 cells were seeded per well in 8-well IbiTreat µ-slides (Ibidi, 80826).

#### Starvation experiments and treatments

For Trp starvation experiments, one day after seeding, cells were washed with PBS (Biotech, Cat# P04-36500) and customized Trp-free DMEM (Gibco, ME15175L1) containing 4.5 g/L glucose, supplemented either with 10% dialyzed FBS (Life Technologies, 26400044), 2 mM L-glutamine (Gibco, 25030-024), 1 mM sodium pyruvate (Gibco, 11360039) and 100 U/mL penicillin and 100 µg/mL streptomycin (Gibco, 15140122) or supplemented with only 3 mM L-glutamine (Gibco, 25030-024) was added to the cells. Trp was dissolved in cell culture grade water (Corning, 25-055-CV) and added fresh at a final concentration of 78 µM or complete DMEM (Gibco, 31053-028) was used as Trp-containing control medium. For the proteome measurements, the cells were cultured in the presence of 78 µM (+Trp), or in the presence of less than 0.4 µM Trp (-Trp). Unless specified otherwise, Trp starvation was performed for 24 h.

For the non-Trp stress proteome experiments, all starvations were performed for 24 h. The following media were utilized: -all aa HBSS (Cat# P04-32505, BioTech), -all aa DMEM (Cat# P04-01507, BioTech); Glucose (Cat# A24940-01, Gibco) was added to reach 4.5 g/L, Met control medium (Cat# 21875-034, Gibco) and Met starvation medium (Cat# A14517-01, Gibco).

For cell treatments, the following compounds were used: EGF (Peprotech, AF-100-15) was diluted in PBS (SERVA, 47302.03) with 0.1% BSA (Carl Roth, 8076.5) and added directly into the media at a final concentration of 10 ng/mL EGF (immunofluorescence experiments) for the indicated time points. Cycloheximide (Sigma-Aldrich, C4859) was diluted in water and directly added to the media at a final concentration of 2 µg/mL for the indicated time points or 5 µg/mL for 24 h. Inhibitors were diluted in DMSO (Sigma-Aldrich, D2650) and cells were treated with 10 µM 4EGI-1 (Tocris, 4800), 10 µM Afatinib (Selleckchem, S1011), 100 nM AZD8055 (MedChem Express, HY-10422), 100 nM Bafilomycin A_1_ (MedChem Express, HY-100558), 10 µM Erlotinib (Selleckchem, S7786), 1 µM Pictilisib (GDC0941, Axon, 1377), and 1 µM StemReginin 1 (SR1, Merck Millipore, 182706) for the indicated time points.

### Generation of transgenic cell lines

#### Transient siRNA-mediated knockdown

Simultaneous siRNA knockdown of all three RAS isoforms (KRAS/HRAS/NRAS) was induced using 10 nM of each KRAS, HRAS and NRAS (total siRNA 30 nM) ON-TARGETplus human SMARTpool siRNA (Dharmacon, L-005069-00-0005, L-004142-00-0005, L-003919-00-0005) for 8 h followed by a medium change. Transfections were preformed using Lipofectamine 3000 (Invitrogen, L3000008) according to the manufacturer’s protocol 48 h (first transfection) before cells were cultured in Trp-containing or Trp-free medium for 24 h.

For siRNA-mediated knockdown of SESN2 ON-TARGETplus human SMARTpool siRNA (Dharmacon, L-019134-02-0005) was used at a final concentration of 40 nM or 60 nM for 8 h followed by a medium change. Transfection was performed with Lipofectamine RNAiMAX (Thermo Fisher Scientific, 13778100) according to the manufactureŕs protocol 24 h before cells were cultured in Trp-containing or Trp starvation medium for 24 h.

ON-TARGETplus non-targeting pool siRNA (Dharmacon, D-001810-10-05) was used for all knockdowns as a control at the same concentration as indicated. Knockdown efficiency was confirmed by gene and protein expression analysis using qRT-PCR and immunoblot.

#### Stable knockdown of AHR

Short hairpin RNA (shRNA) knockdown of AHR in LN-18 cells was performed as previously described in Sadik et al. for U-87 MG cells using the AHR-targeting shERWOOD UltramiR lentiviral shRNA system (transOMIC Technologies, TLHSU1400-196-GVO-TRI) ^61^. In brief, cells were incubated with viral culture supernatants containing either the shAHR or shC (control) sequences for 24 h in the presence of 8 μg/mL polybrene (Merck, TR-1003-G) to facilitate viral infection and subsequent integration of the shRNA sequences. Selection of the transduced cells was performed by first cell sorting using a BD FACSAria fusion (BD Biosciences), selecting only cells expressing ZsGreen. Second, transduced cells were selected by adding 1 μg/mL puromycin (AppliChem, A2856) to the culture media. Stable knockdown of AHR was confirmed at the mRNA and protein levels by qRT-PCR and immunoblot analysis.

#### AHR overexpression

AHR cDNA was amplified by PCR and cloned into the Gateway® pDONR201 vector with primers placed at the end or start positions of the AHR CDS. AttB sites were added to the CDS by a two-step PCR. The first PCR was performed with hybrid primers, consisting of half of the AttB sites and the other half being gene specific. The second PCR was done with primers covering the complete AttB sites.

After sequence verification, the AHR CDS was cloned into the following Gateway® (GW) destination vectors (derived from pDEST26 vectors) with a C-terminal protein-tag: pGW-FLAG, pGW-HA and pGW-MYC by Gateway®-specific LR-reaction following the manufacturer’s protocol (Invitrogen). In addition, the AHR CDS was generated with STOP-codon in the Gateway-frame and cloned into pGW-FLAG with an N-terminal FLAG-tag by Gateway-specific LR reaction. Plasmids were amplified using DB3.1 and Stabl3 E.coli strains and extraction was performed with the QIAprep Spin Miniprep Kit (Qiagen, 27104) and NucleoBond Xtra Midi kit (MACHEREY-NAGEL, 740410.50).

The day after seeding, cells were transiently transfected with Lipofectamine 3000 (Invitrogen, L3000008) following the manufactureŕs protocol with one of the following plasmids pGW-AHR-HA, pGW-AHR-MYC, pGW-AHR-FLAG or pGW-FLAG-AHR, or the respective empty control plasmids pGW-FLAG, pGW-HA and pGW-MYC. 2.5 µg DNA was used per 6-well plate. Media was changed after 6 h of transfection and cells were harvested after 24h. Overexpression was confirmed on mRNA and protein level by qRT-PCR and immunoblot.

#### EIF4EBP1 Ala phospho mutant expression

The coding sequence of human EIF4EBP1 (NM_004095.4) was amplified from cDNA derived from HepG2 cells using primers containing restriction sites using Q5 polymerase (New England Biolabs, M0491). PCR products were cloned into suitable vectors using restriction digest followed by ligation. pCDNA3.1 (Invitrogen, V86020) with an N-terminal Myc-DKK tag and pETDuet-1 (Sigma-Aldrich, 71146) were used for expression in mammalian cells and bacteria, respectively. Mutations T37A T46A were sequentially introduced by PCR using overlapping primers and whole-vector amplification, followed by DpnI (New England Biolabs, R0176S) digestion.

Recombinant H6-EIF4EBP1 was produced in BL21 (DE3) E. coli (Novagen, 69450) co-expressing lambda protein phosphatase and purified with the QIAexpress Ni-NTA Protein Purification System (Qiagen, 30210) according to the manufacturers protocol. We quantified the amount of recombinant H6-EIF4EBP1 (100 ng) using immunoblotting. To ensure the concentration was comparable to endogenous levels, we loaded a cell lysate sample alongside our EIF4EBP1 sample. Furthermore, we validated the concentration by loading a commercial EIF4EBP1 sample with a known concentration for comparison.

Expression of T37A T46A mutated and *wt* EIF4EBP1 was induced using 0.5 µg of the respective plasmid, created as described above. Transfections were preformed utilizing Lipofectamine 3000 (Invitrogen, L3000008) according to the manufacturer’s protocol 48 h before cells were cultured in Trp-containing or Trp-free medium for 24 h.

### Human glioblastoma samples

Tumour specimens of patients diagnosed with glioblastoma (WHO grade IV, IDH wildtype) were obtained from the Institute of Neuropathology, Heidelberg University Hospital, according to the regulations of the Tissue Bank of the National Center for Tumor Diseases (NCT), Heidelberg University Hospital, under the ethics board approval S-318/2022.

### MALDI mass spectrometry imaging (MSI)

#### MALDI-MSI sample preparation

Frozen glioblastoma samples were cut into 10 μm thick sections with a CM1950 cryostat (Leica Biosystems) and mounted onto ITO coated glass slides (Bruker Daltonics, 8237001) for MALDI MSI. Slides were stored in slide boxes (neoLab, 2-3080), covered with foil, vacuumed (CASO) and stored at -80°C until further processing. Consecutive tissue sections were stained with hematoxylin and eosin (HE) and annotation of tumour tissue regions was performed by a clinically experienced neuropathologist. Immediately before matrix coating, the frozen slides were equilibrated at room temperature (RT) and dried for 10 min in a desiccator (SP Bel-Art).

Glioblastoma samples were prepared with the following matrix application protocol: 100 µL of a 5 mg/mL deuterated tryptophan (D5-Trp) solution (Cayman Chemicals, 34829) in ACN/H2O (50:50, v/v) (Honeywell, 34967) was added to 25 mg/mL 2,5-dihydroxybenzoic acid (Alfa Aesar, A11459) in ACN/H2O/TFA (49.4:49.4:0.2, v/v/v) (Merck KGaA, 1.08262.0025) solution and sprayed onto tissue sections with the following parameters: nozzle temperature 75°C, 12 layers, flow rate 0.11 mL/min, velocity 1200, track spacing 2 mm, pattern CC, pressure 10 psi, dry time 0s, nozzle height 40 mm.

#### Magnetic resonance MALDI-MSI data acquisition

Data acquisition was performed on a Fourier-transform ion cyclotron resonance (FT-ICR) magnetic resonance mass spectrometer (MRMS; solariX XR 7T, Bruker Daltonics) in two steps (**Table 1**). First, method 1 optimized for detection of Trp was used at 100 μm step-size. Thereafter, method 2 optimized for detection of ceramides was used on the same tissue section with an XY-offset of 50 μm at step-size 100 μm. Peak filtering was set to SNR > 3 and an absolute intensity threshold of 10^5^ a.u.

**Table 1.** MRMS MALDI acquisition methods for glioblastoma samples.

|  | **Method 1**<br>**optimized for Trp** | **Method 2**<br>**optimized for Ceramides** |
| --- | --- | --- |
| **Online calibration** | D5-Trp 210.128538 | PC (34:1) 760.585082 |
| **Size** | 4M | 2M |
| **Mass range** | m/z 75-800 | m/z 300-800 |
| **Transient length** | 1.47 sec | 2.93 sec |
| **Resolving power at m/z 400** | 200,000 | 390,000 |
| **Laser shots** | 300 | 300 |
| **Frequency** | 2000 Hz | 2000 Hz |
| **Laser focus** | small | small |
| **Q1 Mass** | 200 | 500 |
| **Funnel RF amplitude** | 80 V | 120 V |
| **Time of flight** | 0.5 ms | 0.5 ms |
| **Q1 isolation** | 200 ± 100 | off |

#### Raw data evaluation and visualization

Centroided data was imported into SCiLS Lab 2023a Pro software (Bruker Daltonics), then exported as .imzml file and uploaded to the Metaspace platform (https://metaspace2020.eu/)^95^ for annotation of metabolites. Raw spectra were evaluated in DataAnalysis software (Bruker Daltonics), and the smart formula function was used to generate sum formulas that supported the annotations in METASPACE. Unless stated otherwise, ion images from positive ion mode measurements show protonated adducts ([M+H]^+^) and ion images from negative ion mode measurements show chloride adducts ([M+Cl]^-^).

#### Data Processing

Tissue areas were selected using an in-house-built IT tool and saved as regions of interest (ROI) in the raw data file. Data processing was done in R (version 4.2.1). First, centroided data was loaded, data for every ROIs per tissue sample was extracted using an in-house-built R-tool. The data was loaded as a sparse matrix representation using Matrix package, for faster matrix processing. Next, full width at half maximum (FWHM) was calculated as a function of *m/z* per sample, using the *moleculaR* package ^96^. This was followed by peak binning using the MALDIquant package ^97^, and intensity normalization using *moleculaR* package [Root Means Square (RMS) for ceramides-focused datasets and internal standard (IS; D5-tryptophan *m/z* 210.12854) for tryptophan-focused datasets]. Finally, peak filtering was performed with a minimum frequency set to 0.01.

#### Calculation of molecular probability maps (MPMs) and collective projection probability maps (CPPMs)

Ceramide adduct masses ([M+H]^+^, [M+Na]^+^, and [M+K^+^]) were extracted from the LipidMaps database (www.lipidmaps.org). The data representation was first converted from raw ion intensities into spatial point patterns representations, and then MPMs were calculated per molecule (one ceramide or Trp) of interest (MOI). Subsequently, MPMs for each of the ceramides were standardized and then converted into CPPM representation, as described in Abu Sammour *et al.*, 2021 ^96^. Hotspot areas and contours that indicate significantly increased MOI presence were generated for each tissue sample using the *Spatstat R* package.

#### Calculation of Dice Similarity Coefficient (DSC) values

To calculate overlap between Trp’s MPM hotspot contours with each of the ceramides’ CPPM hotspot contours, Dice Similarity Coefficient (DSC) was calculated as described in Abu Sammour *et al.*, 2021 ^96^.

### RNA isolation, cDNA synthesis and qRT-PCR

For RNA isolation, cells were harvested using RTL buffer containing 10 µL/mL beta-mercaptoethanol (Sigma-Aldrich, M3148) and isolated as recommended in the manufactureŕs protocol of the RNeasy Mini Kit (Qiagen, 74106). DNAse digest step was performed as recommended in the protocol using the RNase free DNAse kit (Qiagen, 79254). RNA concentration and quality were determined by Nanodrop (Thermo Fisher Scientific). Subsequently, cDNA was synthetized using 1 µg RNA and the High Capacity cDNA reverse transcriptase kit (Applied Biosystems, 4368813). Quantitative real-time PCR (qRT-PCR) was performed in a 96-well format using the StepOne Plus Real-Time PCR system (Applied Biosystems) and SYBR Select Master Mix (Thermo Fisher Scientific, 4364346). Expression data was processed using StepOne Software v2.3 (Thermo Fisher Scientific) and analyzed with the 2^-ΔΔCt^ method using *18S rRNA* as reference gene.

For polysome profiling, RNA was isolated using TRI reagent (Zymo, R2050-1-200). In brief, 500 µL of each sucrose fraction was mixed with 0.5 µL TRI reagent (Zymo) and 200 µL chloroform (Fisher Scientific, 10452631), mixed and centrifuged at 13,000 x g for 15 min at 4 °C. After centrifugation, 500 µL of the upper phase was transferred in a new tube containing 1 mL isopropanol (VWR, 84881-320) and 2 µL f 15 mg/mL Glycoblue (Invitrogen, AM9515). RNA was precipitated over night at -20 °C followed by centrifugation at 13,000 x g for 15 min at 4°C. Supernatant was discarded and pellet was washed once with 1 mL ice-cold 70% ethanol (Merck, 818760.2500) by centrifugation at 13,000 x g for 15 min at 4°C. Supernatant was discarded followed by a second centrifugation step at 4°C to collect the remaining ethanol. Ethanol was discarded and RNA pellet was air dried for 5-15 min at 25 °C. RNA was dissolved in 12 µL nuclease-free water. The whole RNA solution was used for reverse transcription as described above. After cDNA synthesis, a qRT-PCR was performed for *AHR* as described above. All qRT-PCR primers used in this study are listed in **Table S4**.

### RNA sequencing

Illumina sequencing libraries were prepared using the TruSeq Stranded mRNA Library Prep Kit (Illumina, 20020595) according to the manufacturer’s protocol. Briefly, poly(A)+ RNA was purified from a maximum of 500 ng of total RNA using oligo(dT) beads, fragmented to a median insert length of 155 bp and converted to cDNA. The ds cDNA fragments were then end-repaired, adenylated on the 3′ end, adapter ligated and amplified with 15 cycles of PCR. The libraries were quantified using Qubit ds DNA HS Assay kit (Life Technologies-Invitrogen, Q33231) and validated on an Agilent 4200 TapeStation System (Agilent technologies, 5067-5582, 5067-5583). Based on Qubit quantification and sizing analysis multiplexed sequencing libraries were normalized, pooled and sequenced using the NovaSeq 6000 Paired-End 100bp S4 flowcell (Illumina, 20028313) with a final concentration of 300 pM spiked with 1% PhiX control (Illumina, 15051973).

#### RNA-seq data processing

We used the DKFZ/ODCF workflows for RNAseq v1.3.0-0, Alignment and QC v1.2.73-3 (https://github.com/DKFZ-ODCF) deployed on the Roddy framework (Roddy v3.5.9; Default-Plugin v1.2.2-0; Base-Plugin v1.2.1-0; https://github.com/TheRoddyWMS/). Paired end FASTQ reads were aligned using the STAR aligner v2.5.3a^98^ by a 2-pass alignment. The reads were aligned to a STAR index generated from the 1000 genomes assembly, gencode 19 gene models (1KGRef_PhiX) and for a sjbdOverhang of 200. The alignment call parameters were -- twopassMode Basic --twopass1readsN -1 --genomeLoad NoSharedMemory --outSAMtype BAM Unsorted SortedByCoordinate --limitBAMsortRAM 100000000000 --outBAMsortingThreadN=1 --outSAMstrandField intronMotif --outSAMunmapped Within KeepPairs --outFilterMultimapNmax 1 --outFilterMismatchNmax 5 --outFilterMismatchNoverLmax 0.3 --chimSegmentMin 15 --chimScoreMin 1 –chimScoreJunctionNonGTAG 0 --chimJunctionOverhangMin 15 --chimSegmentReadGapMax 3 --alignSJstitchMismatchNmax 5 -1 5 5 -- alignIntronMax 1100000 --alignMatesGapMax 1100000 --alignSJDBoverhangMin 3 --alignIntronMin 20. Duplicate marking of the resultant main alignment files, as well as generating BAM indices was done with sambamba v0.6.5 ^99^. Quality control analysis was performed using the samtools flagstat command (samtools v1.6), and the rnaseqc tool^100^.

Featurecounts from the subread package v1.6.5 was used to perform gene specific read counting over exon features based on the gencode 19 gene models^101^. Strand unspecific counting was used. Both reads of a paired fragment were used for counting and the quality threshold was set to 255.

#### Gene expression analysis and gene set testing

The raw RNA-seq counts were imported into R and saved as DGELists^102^. Genes with low counts across all samples were filtered using the function filterByExpr followed by trimmed mean of M values (TMM) normalization^102^ and variance modeling using voom^103^. Batch effects were determined on the principal component analysis (PCA) projections and were corrected by a linear regression model. Differential gene expression was performed using the limma RNA-seq pipeline^103^. Differentially regulated genes were considered significant at a p-value of less than or equal to 0.05. We retrieved the gene sets of the AHR-signature ^61^ and autophagy regulators ^74^ for gene set testing. Comparing the state of activity of any gene set was performed by a non-competitive gene set test using ROAST^104^. Multiple testing correction was performed by applying the Benjamini-Hochberg procedure.

### TCGA glioblastoma expression data

#### Data download

We downloaded and manually curated the metadata entries of 614 submitted glioblastoma patient samples. We excluded 42 entries that were either, duplicates, referring to normal tissue, control analytes, resected from the wrong site or of recurrent tumours. We selected the patient data generated on the two channel Agilent 244K Custom Gene Expression array because it was used for all remaining 572 samples. The Cy3 channel was hybridized with the Stratagene Universal RNA Reference and the Cy5 channel was hybridized with the sample. We used the unique identifiers to download the raw microarray data using GDC-client v1.1.0. (https://portal.gdc.cancer.gov/legacy-archive/search/f)

#### Data annotation

Two different versions of the custom array were used, G4502A-07-1 and G4502A-07-2. Both arrays had ∼87% of common probes, which were later used to merge the patient data from both versions together. The array design files (ADF) and FASTA files were downloaded from https://www.cancer.gov/about-nci/organization/ccg/research/structural-genomics/tcga/using-tcga/technology). We created a new annotation file by aligning the 60 k-mer probes to the non-redundant nucleotide database (https://ftp.ncbi.nlm.nih.gov/blast/db/FASTA/nt.gz; reference build hg38; downloaded on 21.07.2016) by using BLAST+ v2.2.30 and the following call parameters blastn -query unique.probes -task blastn -db nt -out resultblastn.txt -evalue 0.0001 -outfmt “6 std sgi nident staxids sscinames sstitle scomnames sstrand qcovhsp” -num_threads 14. The blast result was annotated using mygene v1.8 and additional gene information was added using the NCBI gene-info file (ftp://ftp.ncbi.nlm.nih.gov/gene/DATA/gene_info.gz; downloaded on 15.08.2016). The annotation file was filtered by removing all hits without the human taxid (9606), and with less than 60 bp matching, a mismatch > 0, and without an “NM_” RefSeq accession prefix.

#### Data processing

We used the Sample and Data Relationship Format files (SDRF) to group the microarray data according to the chip version used. For each group the raw files were imported using the read.maimages function from the limma package. The probes of the raw matrix were background corrected using the “normexp” method with a setoff value of 50, followed by within array normalization using the LOESS-smoothing algorithm. Only probes that were successfully annotated as described above were retained. For every gene, probes were summarized into a single value (gene-set). If a gene was represented by more than three probes, we calculated the median absolute deviation (mad), and if a probe had a value outside the closed interval [-1.5, 1.5], it was counted as an outlier and was filtered out. The remaining probes were averaged to represent the single gene value. All genes with 3 probes or less were averaged, and in the case of genes reported as a single probe, the single probe value was used. The resulting normalized matrix was saved into an MA-list object also including the curated meta- and clinical data. Finally, samples were filtered out if they had a reported IDH mutation, any missing clinical data, or patient age was below 30 years. The final MA-list comprised of 406 patients (GBM406).

#### Feature selection for identifying glioblastoma subgroups

We performed a feature selection step to identify glioblastoma patient subgroups showing high AHR expression and activity, while also reflecting the starvation phenotype observed in the LN- 18 and LN-229 RNA-seq experiments. First, we compiled all differentially expressed genes from the topTables that had an average expression greater than or equal to 1 log2 counts per million, a log2 fold change of 0.58 or higher for upregulated genes and -0.58 or lower for downregulated genes, and adjusted p-value of at least 0.05. The genes fulfilling this criterion in those experiments were 2812 (starvation-features). Next, we estimated immune infiltration scores for the GBM406 patient dataset using the MCP-counter package v1.2.0^105^. Principle component analysis using the FactoMineR package v2.6 was performed with MCP-scores. The starvation-features were correlated using Spearman correlation with the Eigenvalues of each of the first five principal components. Only 1628 genes were left after filtering all other genes that didn’t have a correlation coefficient greater than or equal to 0.3 or less than or equal to -0.3, and a p-value of at least 0.05, with at least one of the first five principal components.

#### Defining glioblastoma subgroups

We applied a graph-based approach to identify glioblastoma subgroups. The subset of the expression matrix comprising the 1628 genes was used for identifying glioblastoma subgroups. First, we created a nearest neighbour graph using the cccd package v1.6. We used the correlation between the genes as a measure of distance, set the k-nearest neighbours to 10, and selected the kd-tree algorithm for the graph embedding^106–108^. We used the Louvain algorithm^109^ for community detection, which defined the seven GB subgroups.

#### Generating enrichment scores

Single sample enrichment scores for the AHR signature^61^ and autophagy regulators^74^ were generated using the GSVA package^110^. In brief, this method accounts for biases resulting from the difference in GC content across genes. Using a Gaussian kernel, the expression values were scaled by estimating the non-parametric kernel of its cumulative density function, which was used for estimating a rank distribution. Kolmogorov Smirnov like random walk statistic was used to calculate a normalized enrichment score based on the absolute difference of the magnitude of the positive and negative random walk deviations.

#### TCGA RPPA data

We downloaded level-4 normalized reverse phase protein arrays (RPPA) of TCGA glioblastoma patients from The Cancer Proteome Atlas (TCPA) (http://tcpaportal.org/tcpa). The data was filtered to include 169 samples, which were in common between both the RPPA and the GBM406 dataset.

#### Interaction network analysis

Volcano plot representation of LN-18 RNA-seq data was performed using tidyverse package in Rstudio. A cut-off (|Log2FC| > 0.5 and adjusted p-value < 0.05) was applied to count significantly altered genes. Autophagy-related genes that were upregulated upon Trp restriction were extracted from Bordi et al^74^. The identified autophagy-related genes were subjected to interaction network analysis using STRING database^111^. Physical interaction networks were generated and subjected to k-means clustering (cluster n = 5) to obtain subnetworks, which were further classified based on the functions of the proteins^74^.

### Promoter binding analysis

The AHR binding motif in the *MAP1LC3B* promotor region was retrieved from the Transcription Factor Target Gene Database (http://tfbsdb.systemsbiology.net/)^112^. Information on chromatin state and regulatory elements were derived from the UCSC browser, including ENCODE histone H3 lysine 27 acetylation (ENCODE histone modification tracks), DNase hypersensitivity cluster information (Integrated Regulation from ENCODE, V3), chromatin segmentation states (Broad ChromHMM) and regulatory element interactions based on GeneHancer (GeneHancer Regulatory Elements and Gene Interactions, V2^113^). The reference genome used was hg19. Binding motifs for AHR (*MAP1LC3B* promoter) were visualized in conjunction with chromatin state and interaction data from the UCSC browser.

### Protein isolation and immunoblot

For protein harvest, cells were washed once with ice-cold PBS (Gibco, 14190169) and lysed with radio immunoprecipitation assay (RIPA) buffer containing 1% IGEPAL CA-630 (Sigma-Aldrich, I8896), 0.1% SDS (Carl Roth, 8029.3), and 0.5% sodium deoxycholate (AppliChem, A1531) in PBS supplemented with Phosphatase Inhibitor Cocktail 2 and Cocktail 3 (Sigma-Aldrich, P5726, P0044) and Complete Protease Inhibitor Cocktail (Roche, 11836145001) and centrifuged for 10 min at 13,000 g and 4°C. Protein concentration was determined using Protein Assay Dye Reagent Concentrate (Bio-Rad, 5000006), and absorbance was measured at 595 nm using a spectrophotometer (GE Healthcare). All samples within one experiment were adjusted to the lowest absorbance value. Cell lysates were mixed with 5x Laemmli buffer containing 10% glycerol (Sigma-Aldrich, 15523), 1% beta-mercaptoethanol (Sigma-Aldrich, M3148), 1.7% SDS (Carl Roth, 8029.3), 62.5 mM TRIS base (Sigma-Aldrich, T1503) [pH 6.8], and bromophenol blue (Sigma-Aldrich, B5525), and boiled for 5 min at 95°C. Separation of proteins was performed with SDS polyacrylamide gel electrophoresis (PAGE) using gels with a concentration of 8%, 10%, 14% or 15% acrylamide (Carl Roth, 3029.1) in a Mini-PROTEAN Tetra Vertical Electrophoresis Cell system (Bio-Rad, 1658029FC) with running buffer containing 0.2 M glycine (Sigma-Aldrich, 33226), 25 mM TRIS base (Sigma-Aldrich, T1503), and 0.1% SDS (Carl Roth, 8029.3) at 80 – 150 V. Proteins were blotted onto a PVDF membrane (Merck Millipore, IPVH00010) or nitrocellulose membrane (Sigma-Aldrich, GE10600001) at 45 V for 2 h using the Mini-PROTEAN Tetra Vertical Electrophoresis Cell System (Bio-Rad, 1658029FC) and the blotting buffer containing 0.1 M glycine (Sigma-Aldrich, 33226), 50 mM TRIS base (Sigma-Aldrich, T1503), 0.01% SDS (Carl Roth, 8029.3) [pH 8.3], and 10% methanol (Merck, 1.06009.2511). Membranes were blocked for 1 h at RT in 5% BSA (Carl Roth, 8076.5) in Tris-buffered saline tween (TBST) buffer (0.15 M NaCl (Sigma-Aldrich, S7653), 60 mM TRIS base (Sigma-Aldrich, T1503), 3 mM KCl (Sigma-Aldrich, P405), and 0.1% Tween-20 (Sigma-Aldrich, P9416), [pH 7.4]). Primary antibodies were diluted as recommended by the manufacturer in 5% BSA in TBST and incubated overnight at 4°C. On the next day, membranes were washed three times for 10 min in TBST buffer and subsequently incubated for 2 h with the respective horseradish peroxidase (HRP)-coupled secondary antibody dissolved in 5% BSA in TBST buffer. After another three 10 min wash steps in TBST buffer, proteins were detected using ECL Western Blotting Substrate (Thermo Fisher Scientific, 32106, Amersham, RPN2235), or SuperSignal West FEMTO (Thermo Fisher Scientific, 34096) under a ChemiDoc XRS+ camera system (Bio-Rad, 1708265) or a Fusion Fx camera (Vilber). Images taken with the ChemiDoc XRS+ were quantified with the Image Lab software (Bio-Rad, v6.0.1). Images taken with the Fusion FX camera were quantified with the ImageQuant TL 1D (v8.2.0). Normalization was performed as described^12^. In brief, the images were first normalized by the pixel values of a single lane to the average value of all lanes in a blot for each antibody. Subsequently, the internally normalized proteins were normalized to the loading control glycerinaldehyd-3-phosphat-dehydrogenase (GAPDH), tubulin (TUBA1B), or vinculin (VCL), as indicated.

Upon separation by SDS-PAGE, EIF4EBP1 runs in three discernible bands (alpha, beta, gamma), all of which can be phosphorylated at T37/46 by MTORC1^114,115^. We therefore quantified across all EIF4EBP1-pT37/46 signals.

### Puromycin Assay

Protein synthesis was measured by puromycin assay. Therefore, 5 µg/mL puromycin (Sigma-Aldrich, P8833) was added directly to the media 5 min prior lysis. Puromycin incorporation was detected by immunoblot analysis, as described above. The entire lane was used for quantification of the puromycin blots.

### Cap pull down

The cells were washed two times in ice-cold PBS and harvested in 1 mL per 15 cm plate of cap pull down lysis buffer (40 mM HEPES, 120 mM NaCl, [pH 7.5], 0.3% CHAPS) supplemented with Phosphatase Inhibitor Cocktail 2 and Cocktail 3 (Sigma-Aldrich, P5726 and P0044) and Complete Protease Inhibitor Cocktail (Roche, 11836145001). The lysed cells of every condition were pooled together (six dishes per condition were seeded). Afterwards, the samples were end over end rotated at 4°C for 20 min and then centrifuged for 3 min at 600 g and 4°C. The protein concentrations were measured in the supernatants using Protein Assay Dye Reagent Concentrate and all samples were adjusted to the lowest value. For the input analysis, 160 µL per condition were mixed with 40 µL of 5x Laemmli buffer [10% glycerol (Sigma-Aldrich, 15523), 1% beta-mercaptoethanol (Sigma-Aldrich, M3148), 1.7% SDS (Carl Roth, 8029.3), 62.5 mM TRIS base (Sigma-Aldrich, T1503) [pH 6.8], and bromophenol blue (Sigma-Aldrich, B5525)], incubated at 95°C for 5 min, vortexed, centrifuged and stored at -20°C. Cap pull down beads (Immobilized γ-Aminophenyl-m7GTP, Jena Bioscience, AC-155) and mock beads (Jena Bioscience, AC-001) were washed twice with lysis buffer. The samples were respectively split into halves. 17.5 µL of cap pull down beads / lysate were added to one half of a sample, 17.5 µL of mock beads / lysate were added to the other half of a sample. The sample-beads mixtures were end over end rotated at 4°C for two hours and then centrifuged at 500 g for 1 min. The supernatants were removed and the pellets resuspended in 500 µL of lysis buffer. After transferring the resuspended pellets into new tubes, they were washed three times with lysis buffer by inverting the tubes six times and centrifugation at 500 g for 1 min. After the last washing step, the buffer was carefully removed and the pellets were dissolved in 65 µL of 1x Laemmli buffer. The samples were gently vortexed and incubated for 10 min at 95°C before freezing them at -20°C.

### Ras pull down assay

For Ras pull down, cells were collected in MLB buffer (RAS Activation Assay Kit, Merck Millipore, 17-218) after 30 min of 10 ng/mL EGF (Preprotech, AF-100-15) stimulation. Protein concentration was determined using BCA assay (Thermo Fisher Scientific, 23227) and pellets were frozen in -80°C. The RAS GTP pull-down assay was performed as described in Heberle et al, 2019^44^. In short, 500 µL protein extracts (800 µg -1 mg, adjusted depending on the lowest concentration in each replicate) were incubated for 45 min at 4°C with 10 µL agarose beads using a RAS-GTP pull-down assay kit (RAS Activation Assay Kit, Merck Millipore, 17-218). Supernatant was recovered after centrifugation, mixed with 40 µL of 1x Laemmli buffer, incubated at 95°C for 5 min, centrifuged and stored at -20°C. For immunoblot analysis of RAS-GTP levels 20 µL of protein extract was separated by gel electrophoresis, blotted and incubated overnight in 5% skim milk (GERBU Biotechnik, 70166) in TBST at 4°C with an anti-RAS antibody (Millipore, 05-516). As a loading control, glutathione-S transferase (GST; CST, 2622) was tested in 5% skim milk in TBST for 2 h at RT. Immunoblots were quantified using ImageJ v.153k. Single lane chemiluminescence values were normalized to the average value of all lanes in a blot for each antibody, and subsequently normalized to the internal loading control GST.

### Kinase assay

The kinase assays were developed based on previous work^116,117^. Media exchange was performed once 10 cm dish (TPP, 93100) was confluent. After treatment, the cells were washed three times in ice-cold PBS and then harvested in 600 µL per plate CHAPS-based IP lysis buffer [40 mM HEPES (Gibco, 15630056), 120 mM NaCl (Sigma-Aldrich, S7653), [pH 7.5] and 0.3% CHAPS (Merck, 331717-45-4)] supplemented with 500 nM Benzamidine (Benzamidine, B6506-5G) and 20 µg/mL Heparin (Sigma-Aldrich, H3149-25KU). The lysate was incubated under gentle agitation for 20 min at 4°C, centrifuged for 3 min at 600 g at 4°C, the pellet was discarded and the supernatant was transferred to fresh tubes. In case of multiple samples, the protein concentration was measured using Protein Assay Dye Reagent Concentrate (Bio-Rad, 5000006) and all samples were adjusted to the lowest value. The lysates were pre-incubated with 10 μL pre-washed Protein G covered Dynabeads (Life Technologies, 10009D) per mL of lysate for 30 min at 4°C under gentle agitation. The pre-cleaned lysates were subdivided, and anti-RPTOR antibodies (Helmholtz Center Munich, 20D12) or isotype control IgG antibodies (Helmholtz Center Munich, 7H8) were added using 7.5 μg antibody per mL of pre-cleaned lysate. Isotype control IgG antibodies (mock antibodies) were used in the same concentration as the protein-specific antibodies. After 30 min at 4°C under gentle agitation, 37.5 μL pre-washed Protein G covered Dynabeads / mL lysate were added, and the incubation was continued for 90 min at 4°C under gentle agitation. Next, beads were washed with CHAPS lysis buffer three times shortly and three times for 10 min at 4°C under gentle agitation.

Following the last wash step with CHAPS lysis buffer, the beads were subdivided and excess liquid was removed. The kinase assays were performed in a final volume of 30 µL, containing kinase assay buffer (final concentration: 40 mM HEPES, 120 mM NaCl, [pH 7.5] and 0.3% CHAPS, 4 mM MnCl2 (Merck, 1059270100), 10 mM DTT (Sigma-Aldrich, D0631), supplemented with 1x Protease inhibitor cocktail without EDTA and 2 µg/mL Heparin), 100 ng recombinant H6-EIF4EBP1 (cloned and purified as described above), 500 mM AZD8055 (MedChem Express, HY-50706) and 0.133 mM ATP (Merck, 74804-12-9).

First, 24.5 µL kinase assay buffer was added to each condition, before 0.5 µL of dried AZD8055, dissolved in kinase assay buffer, or 0.5 µL of kinase assay buffer was added and pre-incubated for 15 min at 4°C before initiation of the kinase reaction. The kinase reactions were started by adding 1 μL of recombinant H6-EIF4EBP1, or kinase assay buffer, and addition of 4 µL 1 mM ATP. The reactions were incubated at 30 °C for the indicated time points, and stopped by the addition of 30 µL 2x Laemmli buffer. Samples were heated for 5 min at 95°C and separated by SDS-PAGE. The MTORC1-mediated phosphorylation of EIF4EBP1-pT37/46 and MTOR levels were run on gradient gels and detected by immunoblotting with specific antibodies. Signals were quantified using ImageQuant (v8.2.0) and are shown as the EIF4EBP1-pT37/46/MTOR ratio.

### Immunofluorescence

For EGFR immunofluorescence experiments, cells were washed with ice-cold PBS (Gibco, 14190169), fixed in 100% methanol (VWR, 85681-320) for 10 min at RT and permeabilized with 0.3% Triton X-100 (Sigma-Aldrich, T8787) in TBS [0.15 M NaCl (Sigma-Aldrich, S7653), 60 mM TRIS base (Sigma-Aldrich, T1503), and 3 mM KCl (Sigma-Aldrich, P405) [pH 7.4]] for 10 min at 37°C. Prior to immunofluorescence, blocking was performed in TBS + 1% BSA (Carl Roth, 8076.5) for 2 h and incubated with anti-EGFR (CST, 4267) and anti-LAMP2 (DSHB, H4B4) antibodies for 3 h at RT. After three wash steps in TBST [0.15 M NaCl (Sigma-Aldrich, S7653), 60 mM TRIS base (Sigma-Aldrich, T1503), 3 mM KCl (Sigma-Aldrich, P405), and 0.1% Tween-20 (Sigma-Aldrich, P9416), [pH 7.4]], anti-rabbit Alexa-488 (A-11008, Invitrogen) and anti-mouse Alexa-647 antibodies (A-32728, Invitrogen) were added for 2 h at RT in the dark. Finally, nuclei were counterstained with 5 µg/mL DAPI (BD Biosciences, 564907) in TBS for 1 min. Microscopy was performed using a CQ1 Confocal Quantitative Image Cytometer (Yokogawa Electric). For nuclear, EGFR focus and counting, binary masks were generated from intensity-thresholded images. For LAMP2 compartment size measurement, images were thresholded using an IJ_Isodata algorithm. The total measured area was normalized to nuclear count per image to determine the mean LAMP2 compartment size per cell. Image analysis was performed using ImageJ v.153k.

For p62 autophagy immunofluorescence experiments, the cells were washed twice with ice cold PBS (Gibco, 14190169) and fixed with 4% PFA (AppliChem, A3813) in PBS for 20 min. Afterwards, the cells were washed three times with PBS before permeabilizing them with 0.1% Triton X-100 (Sigma-Aldrich, T8787) in PBS for 60 seconds. After another three washing steps with PBS, blocking solution (5% FBS and 0.05% Tween20 in PBS) was added to the cells for 60 min. The p62 primary antibody (Progen, GP62-C) was diluted in blocking solution, and the cells incubated in the diluted primary antibody overnight at 4°C in a humid chamber. The next day, the cells were washed three times short and two times for 10 min with PBS. The Alexa Fluor 568 labelled secondary antibody (Invitrogen, A-11075) and Hoechst 33342 (Invitrogen, H3570) were diluted in blocking solution, and added to the cells for 60 min at RT in a dark humid chamber. The cells were washed three times with PBS and then twice with deionized water, before mounting the coverslips with Mowiol® 4-88 with DABCO (1,4-diazabicyclo[2.2.2]octane) and 10% n-propyl-gallate (NPG). For image collection, the Axiocam 702 mono camera was utilized, acquiring seven stacks (0.5 µM thickness) per image. The exposure time was adjusted to the condition with the strongest signal and kept constant throughout all conditions. For quantifying the p62 foci per cell, CellProfiler 4.2.1 was used.

### Macropinocytosis Assay

For the uptake assay, 20 µL of medium with 70 kDa-Dextran Oregon Green (dextran) (Invitrogen, D7173) with a final concentration of 0.1 mg/mL and 10 ng/mL EGF were added for 30 min. Next, cells were washed twice with ice-cold PBS and fixed with 4% formaldehyde (AppliChem, A3813) in PBS for 20 min at RT. Fixed cells were washed with PBS and incubated with 10 mg/mL DAPI (Serva Electrophoresis, 18860) in PBS for 10 min. Finally, cells were washed again with PBS and imaged using an AxioObserver.Z1, equipped with an LSM780 ConfoCor 3 microscope with a 63x / 1.4 Oil DIC M27 Plan-Apochromat objective and ZEN 2012 (Zeiss, black edition, v8,1,0,484) software. Nuclear staining using DAPI was imaged with an UV diode (405 nm) and the dextran detection (488 nm) was performed using an argon multiline (458/488/514 nm). Detector gain and detector offset were adjusted once and never changed for an entire dataset. Raw images (CZI files) were subjected for further analyses in Fiji.

Dextran fluorescence was analyzed with Fiji version 1.52p using a background subtraction of 3, a Gaussian Blur filter of 1, threshold adjustment from 3500-max, a prominence of 10, and the ‘Analyze Particles’ function with a particle size from 5-infinity. The number (‘count’) of macropinosomes was then divided by the number of respective cells displayed in the DAPI channel in the analysed microscopy picture. The number of macropinosomes per cell were compared between at least 5 independent fields of view from 4 independent datasets. For presentation in figures, ZEN 3.0 (Zeiss, blue edition) was used, and representative regions of interest for each condition were exported as TIFF with no compression using the ZEN ‘Best Fit’ option. Dextran green fluorescence was pseudo-coloured white. Finally, brightness or contrast were adjusted for better visibility.

### Lysotracker

For lysosome tracking, 20 min before live cell imaging, cells were washed with PBS and 10 nM LysoTracker™ Red DND-99 (lysotracker) (Invitrogen, L7528) and 10 mg/mL DAPI (Serva Electrophoresis, 18860) in PBS were added. Living cells were imaged as above with an AxioObserver.Z1, equipped with an LSM780 ConfoCor 3 microscope with a 63x / 1.4 Oil DIC M27 Plan-Apochromat objective and ZEN 2012 (black edition, v8,1,0,484) software. Nuclear staining using DAPI was imaged with an UV diode (405 nm) and lysotracker with a 561 nm laser. Detector gain and detector offset were adjusted once and never changed for an entire dataset. Raw images (CZI files) were subjected for further analyses in Fiji. Lysotracker was analyzed with Fiji version 1.52p using a background subtraction of 3, a Gaussian Blur filter of 1, threshold adjustment from 3500-max, a prominence of 10, and the ‘Analyze Particles’ function with a particle size from 15-infinity. The raw integrated density (RawIntDen) value was then divided by the number of respective cells displayed in the DAPI channel in the analysed microscopy picture. The intensity of lysotracker foci per cell was then compared across at least 3 independent fields of view from 5 independent datasets.

### Simultaneous proteo-metabolome liquid-liquid extraction and measurement

Proteome extraction from cells was done by a simultaneous proteo-metabolome liquid-liquid extraction ^118^. The cells were washed three times with PBS and cell metabolism was quenched by addition of 500 µL ice-cold methanol (Fisher Chemical, 10653963) and 500 µL MS-grade water (Millipore, Direct Water Purification System). Lysates were scraped and transferred to tubes followed by the addition of 500 µL chloroform. After agitation in a cell shaker at 4°C for 20 min and 500 rpm, phase separation was performed by centrifugation at 4°C for 5 min at 16,100 g. Subsequently, after removing the liquid polar and non-polar phases, the solid interphases containing the proteomes were washed with methanol. Finally, interphases were dried, covered with 50 µL methanol, and stored at -80°C until further processing.

#### Protein extraction from interphases

To extract proteins, 60 µL of 8 M urea (Sigma-Aldrich, 51456) in 100 mM ammonium bicarbonate (Sigma-Aldrich, 09830-500G), pH 8.2 were added to the interphases followed by 240 µL of 100 mM NH_4_HCO_3_, pH 8.2. To bring proteins into solution, samples were sonicated with a tip sonicator (Thermo Fisher Scientific, 10588013). Protein concentration was determined using a microplate BCA protein assay kit (Thermo-Fisher-Scientific, 23227) following the manufacturer’s instructions. For protein determination, samples were diluted 1:50 in MilliQ water. A BSA standard was used to calibrate the assay across the concentration range of 0 - 200 µg/mL. The absorbance was measured at 580 nm using a plate reader (BMG Labtech, PHERAstar FSX). Extracts from samples that had been cultured with 0.4, 0.2 and 0 µM Trp were pooled.

#### Digestion and desalting

100 µg of dissolved protein was transferred into a new vial and filled to a final volume of 100 µL with the extraction buffer. Samples were incubated with 1 M DTT (Sigma-Aldrich, D0631) in 0.1 M triethylammonium bicarbonate (TEAB) (Sigma-Aldrich, 15715-58-9) to a final concentration of 10 mM DTT on a shaker for 30 min at 55°C and 800 rpm. Afterwards, alkylation was performed by 0.5 M iodoacetamide (IAA) (Sigma-Aldrich, I1149). IAA was added to a final concentration of 20 mM and incubated in the dark for 30 min. To quench the remaining IAA, DTT (1 M DTT in 0.1 M TEAB) was added. Digestion of the proteins was performed by the addition of trypsin (Gibco, 15400054) in a trypsin:protein ratio of 1:20. After overnight digestion at 37°C, the reaction was stopped by adding 100% formic acid (FA) (Fisher Scientific, 10596814) to achieve a final concentration of 1% FA in each sample. Afterwards, peptide samples were desalted using Oasis HLB 1 cc Vac Cartridge (Waters, 186000383). For this, the cartridges were first activated with 1 mL of 100% methanol, followed by 1 mL of 95% ACN (Fisher Scientific, 10616653), 0.1% FA. Next, equilibration was performed by adding twice 1 mL of 0.1% FA. Peptide samples were slowly loaded onto the cartridge in 1 mL 0.1% FA. After washing twice with 1 mL 0.1% FA, samples were eluted from the cartridge with 1 mL 70% ACN, 0.1% FA. Samples were dried in a SpeedVac (Eppendorf, Concentrator 5301) and dried peptides were stored at -80°C until further processing.

#### LC-MS/MS analysis

For LC-MS/MS analysis of the Trp stress proteome, the dried tryptic peptides were dissolved in 20 µL 0.1% FA. The samples were injected on a nano-ultra pressure liquid chromatography system (Dionex UltiMate 3000 RSLCnano pro flow, Thermo Fisher Scientific) coupled via an electrospray ionization (ESI) source to an orbitrap hybrid mass spectrometer (QExactive, Thermo Scientific). The samples were loaded (5 µL/min) on a trapping column (nanoE MZ Sym C18, 5 μm, 180 µm x 20 mm, Waters; buffer A: 0.1% FA in HPLC-H_2_O; buffer B: 100% ACN, 0.1% FA) with 100% buffer A. After sample loading, the trapping column was washed for 5 min with 100% buffer A (5 μL/min) and the peptides were eluted (300 nL/min) onto the separation column (nanoE MZ PST CSH, 130 A, C18, 1.7 μm, 75 μm x 250 mm, Waters) and separated with a gradient of 2−30% B in 60 min. The spray was generated from a steel emitter (Fisher Scientific) at a capillary voltage of 1850 V. MS/MS measurements were carried out in data-dependent acquisition mode (DDA) using a normalized HCD collision energy of 25% and a loop count of 15. MS scan was performed over an m/z range from 400-1200, with a resolution of 70,000 at m/z 200 (maximum injection time = 240 ms, AGC target = 1e6). MS/MS spectra were recorded over a m/z range of 100-2000 m/z with a resolution of 17,500 at m/z 200 (maximum injection time = 50 ms, maximum AGC target = 1e5, intensity threshold: 5e3), a quadrupole isolation width of 2 Da and an exclusion time of 20 seconds.

For LC-MS/MS analysis of the -all aa DMEM, -all aa HBSS and Met stress proteomes, the dried tryptic peptides were dissolved in 20 µL 0.1% FA. Samples were injected on a nano-ultra pressure liquid chromatography system (Vanquish Neo UHPLC System, Thermo Fisher Scientific) coupled via an electrospray ionization (ESI) source to an Orbitrap Eclipse (Thermo Scientific). The samples were loaded (60 µL/min) on a trapping column (nanoE MZ Sym C18, 5 μm, 180 µm x 20 mm, Waters; buffer A: 0.1% FA in HPLC-H_2_O; buffer B: 80% ACN, 0.1% FA) with 100% buffer A. After sample loading, the trapping column was washed and the peptides were eluted (300 nL/min) onto the separation column (nanoE MZ PST CSH, 130 A, C18 1.7μ, 75 μm x 250 mm, Waters) and separated with a gradient of 1-40% B in 90 min. The spray was generated from a steel emitter (Fisher Scientific) at a capillary voltage of 1850 V. MS/MS measurements were carried out in data-dependent acquisition mode (DDA) using a normalized HCD collision energy of 30% and a cycle time of 3s. The MS scan was performed over an m/z range from 375-1500, with a resolution of 240,000 at m/z 200 (maximum injection time= 50 ms, AGC target= 4e5). MS/MS spectra were recorded over a m/z range of 135-2000 m/z (maximum injection time= 35 ms, maximum AGC target= 1e5, intensity threshold: 5e3), a quadrupole isolation width of 0.8 Da and an exclusion time of 60 seconds in the ion trap.

#### LC-MS/MS data processing

LC-MS/MS raw files were analysed with ProteomeDiscoverer 2.4 (Thermo Fisher Scientific). For peptide and protein identification, the LC-MS/MS were searched with SequesHT against a human database (SwissProt, 20,369 entries) and a contaminant database (116 entries). The following parameters were used for the data-base search: mass tolerance MS1: 10 ppm, mass tolerance MS2: 0.02 Da for MS/MS analysis in the orbitrap and 0.5 for MS/MS analysis in the ion trap, fixed modification: carbamidomethylation (cysteine), variable modification: Oxidation (Methionine), variable modification at protein N-terminus: Acetylation, Methionine loss, Methionine loss + Acetylation. Trp-Phe exchanges were included as a variable modification for analysis of the Trp stress proteome. Percolator were used for FDR calculation. For feature detection, Minora Feature Detection was used with default settings. For label-free quantification, the Precursor Ions Quantifier was used with the following parameters: Peptides to use: unique peptides, Precursor Abundance Based On: Area, Minimum Replicate Features: 100% for the Trp stress proteome and 75% for generalized amino acid stress (HBSS, DMEM) and methionine stress proteomes, Normalization Mode: Total Peptide Amount, Protein Abundance Calculation: Summed Abundances, Top N: 3, Hypothesis testing: t-test (Background Based). Adjusted p-values were calculated using Benjamini–Hochberg correction. Venn diagrams show proteins up- or downregulated in different conditions. Proteins are defined as regulated if they have a fold change of at least 1.5 with an adjusted p-value of lower than 0.05.

GO enrichment was performed with g:profiler^119^. Resulting p-values were corrected with the Benjamini-Hochberg method. Visualization of results was done using the ggplot2 package in R^120^. To assess if protein synthesis was disrupted at tryptophan positions leading to shorter proteins, the protein coverage by mass spectrometry was analyzed. For this, proteins that were upregulated in low tryptophan conditions were considered. A density plot was created showing the distribution of peptides of proteins upregulated under Trp starvation. Additionally, a heatmap was created showing the peptide coverage for all regulated proteins. Peptides were considered as „quantified“ if they were assigned by Proteome Discoverer with at least confidence „High“ in any of the low tryptophan samples.

#### Extraction of intracellular Trp and quantification by mixed mode reversed phase-anion exchange UPLC-MS/MS

The cells were treated as described in the Simultaneous proteo-metabolome liquid-liquid extraction paragraph. A fully ^13^C, ^15^N labelled amino acid standard (Cambridge Isotope Laboratories, MSK-CAA-1) was spiked into samples at the first step of the extraction. Dried polar phases obtained from simultaneous extraction were dissolved in 100 µL of water containing 5 mM ammonium formate (NH_4_FA) (Sigma-Aldrich, 70221-100G-F) and 0.15% FA (Fisher Scientific, 10596814). 1 µL of each sample was injected. Analytes were separated at 40°C on an Atlantis Premier BEH C18 AX column (1.7 µm, 2.1 x 150 mm, Waters, 186009361) using an Acquity Premier UPLC system (Waters).

A gradient was run at a flowrate of 0.3 mL/min with mobile phase A (5 mM NH_4_FA and 0.15% FA in water) and mobile phase B (10 mM NH_4_FA and 0.15% FA 80% ACN) as follows: 5% B to 15% B in 2 min, 15% B to 70% B in 1.5 min, 70% B to 95% B in 0.5 min followed by 1 min of elution at 95% B and re-equilibration of the column to initial conditions over 2 min. Trp was detected using a Xevo-TQ XS Mass spectrometer (Waters) equipped with an electrospray ionization source running in positive mode. The transition from 205.1 -> 146.2 for endogenous Trp and 218.1 -> 156.1 were used for quantification. The cone voltage was set to 14 V and the collision energy was set to 18 V. Raw files were analysed in TargetLynx (Waters, V4.2 SCN1012). Resulting peak areas of endogenous and ^13^C, ^15^N tryptophan were further analysed in R and resulting tryptophan concentrations were normalised to cell numbers.

For the determination of the intracellular Trp concentration in µM, the average cell size (11.4 µm) was determined as the size per cell via the CytoSMART Exact cell counter (Axion BioSystems) with a lower size gate of 8 µm. Assuming a sphere, the cell volume was estimated to be 775.7 µm^3^.

#### Extraction of sphingolipids

For the measurement of sphingolipids in LN-18 cells, the cells were washed with 5 mL PBS (4°C) and trypsinized with 1 mL 0.25% Trypsin-EDTA per dish (Gibco, 25200-056) for 5 min at 37°C and 5% CO2. After culture medium (4 mL) has been added, cells were pelleted (500 g, 5 min, 4°C), washed twice with ice-cold PBS (1.0 mL and 0.5 mL, 4°C), centrifuged (3000 g, 5 min, 4°C), frozen in liquid nitrogen, and stored at -80°C.

Sphingolipids were extracted from LN-18 cell pellets by successive addition of PBS pH 7.4, methanol, chloroform, and saline to a final ratio of 14:34:35:17^121,122^. The organic phase was evaporated to dryness using an Eppendorf Concentrator Plus System (Eppendorf, 5305000509; high vapor pressure application mode), and the remaining lipid film was dissolved in methanol, centrifuged twice at 21,100×g, 4°C for 5 min, and subjected to UPLC-MS/MS analysis. Internal standards used (Sigma-Aldrich): D-erythro-sphingosine-d7, N-heptadecanoyl-D-erythro-sphingosine, D-glucosyl-β-1,1′-N-heptadecanoyl-D-erythrosphingosine, N-lauroyl-ceramide-1-phosphate, N-heptadecanoyl-D-erythro-sphingosylphosphorylcholine, and D-erythro-sphingosine-d7-1-phosphate.

#### Analysis of sphingolipids by reversed phase UPLC-MS/MS

Chromatographic separation of sphingosines (Sph), (dihydro)ceramides ([dh]Cer), hexosyl-ceramides (HexCer), ceramide-1-phosphates (C1P), and (dihydro)sphingomyelines ([dh]SM) was carried out at 45°C on an Acquity UPLC BEH C8 column (130Å, 1.7 μm, 2.1 × 100 mm, Waters, 186002878) using an ExionLC AD UHPLC system (Sciex). The gradient of mobile phase A (water/ACN, 90/10, 2 mM ammonium acetate) and mobile phase B (ACN/water, 95/5, 2 mM ammonium acetate) was ramped at a flow rate of 0.75 mL/min from 75% to 85% B within 5 min and to 100% B within another 2 min, followed by 13 min of isocratic elution.

Sphingolipids were analyzed in the positive ion mode by scheduled multiple reaction monitoring (MRM) using a QTRAP 6500^+^ Mass Spectrometer (Sciex), which was equipped with an electrospray ionization source. Transitions from [M+H]^+^ to [M+H-H_2_O]^+^ (Sph, dhCer), m/z = 184.1 ([dh]SM), and m/z = 264.4 (Cer, HexCer, C1P) were selected for quantitation. The curtain gas was set to 40 psi, the collision gas to medium, the ion spray voltage to 5000 V, the heated capillary temperature to 500°C, and the sheath and auxiliary gas pressure to 40 psi. The declustering potential was adjusted to 30 V (Sph, [dh]Cer, C1P) or 40 V (HexCer, [dh]SM), the entrance potential to 5 V (HexCer) or 10 V (Sph, [dh]Cer, C1P, [dh]SM), the collision energy to 20 eV (Sph), 30 eV ([dh]SM), 40 eV ([dh]Cer, C1P), or 50 eV (HexCer), and the collision cell exit potential to 5 V (C1P), 10 V ([dh]SM), 20 V ([dh]Cer, HexCer), or 25 V (Sph).

In variation to the procedure described above, sphingosine-1-phosphate (S1P) was separated on an Acquity UPLC CSH C18 column (130Å, 1.7 μm, 2.1 × 50 mm, Waters, 186005296) at 55°C. The LC system was operated at a flow rate of 0.55 mL/min using water/ACN (80/20) with 0.1% formic acid as mobile phase A and isopropanol/ACN (80/20) with 0.1% formic acid as mobile phase B. Initial conditions (60% B) were kept for 3 min, linearly increased to 70% B within 2 min and further to 100% B within 0.4 min, followed by isocratic elution for 1.6 min.

For the analysis of S1P in the positive ion mode ([M+H]^+^) by MRM, [M+H-H_3_PO_4_-H_2_O]^+^ (m/z = 264.2) was detected as fragment ion. The curtain gas was set to 40 psi, the collision gas to low, the ion spray voltage to 4500 V, the heated capillary temperature to 500°C, the sheath gas pressure to 60 psi, the auxiliary gas pressure to 30 psi, the declustering potential to 40 V, the entrance potential to 10 V, the collision energy to 20 eV, and the collision cell exit potential to 20 V. Relative proportions of total ceramides (calculated as sum of ceramide species analyzed) are given as percentage of the sum of all sphingolipids determined in the corresponding sample (= 100%). Mass spectra were acquired and processed using Analyst 1.7.1 or 1.7.2 (Sciex) and Analyst 1.6.3 (Sciex), respectively.

#### Targeted Proteomics samples preparation

Samples were extracted via the simultaneous proteo-metabolome liquid-liquid extraction as described above. The dried interphases were solubilized in 50 µL 1% sodium deoxycholate in 50 mM ammonium bicarbonate buffer pH 8.5. Samples were heated at 95°C for 5 min and sonicated in bath sonicator for 10 min. Protein concentration was measured by BCA assay (Thermo Fischer Scientific) and 30 µg proteins per samples were spiked with synthetic heavy-labelled peptides (**Table 2**). Synthetic peptides were C-terminally labeled with heavy lysine (^13^C_6_, ^15^N_2_) or arginine (^13^C_6_, ^15^N_4_) residues and spiked to reach an amount of approx. 12 or 120 fmol on column depending on the previously measured peptide signals. Samples were reduced with 1 mM DTT for 30 min at RT and alkylated for 30 min with 5.5 mM IAA for 30 min in the dark. Trypsin (Promega) was added to a 1/100 trypsin to protein ratio and the samples were incubated overnight at 37°C. Samples were acidified to 2% TFA and vortexed to precipitate the deoxycholate. Samples were spun and the supernatant was desalted on reverse-phase S cartridges (Agilent) using an Agilent Bravo automated liquid handling platform (Agilent). Samples were lyophilized overnight and resuspended in 0.1% formic acid in water. Peptide concentrations were measured on a NanoDrop (Thermo Fischer Scientific) and adjusted to 100 ng/µL.

**Table 2.**
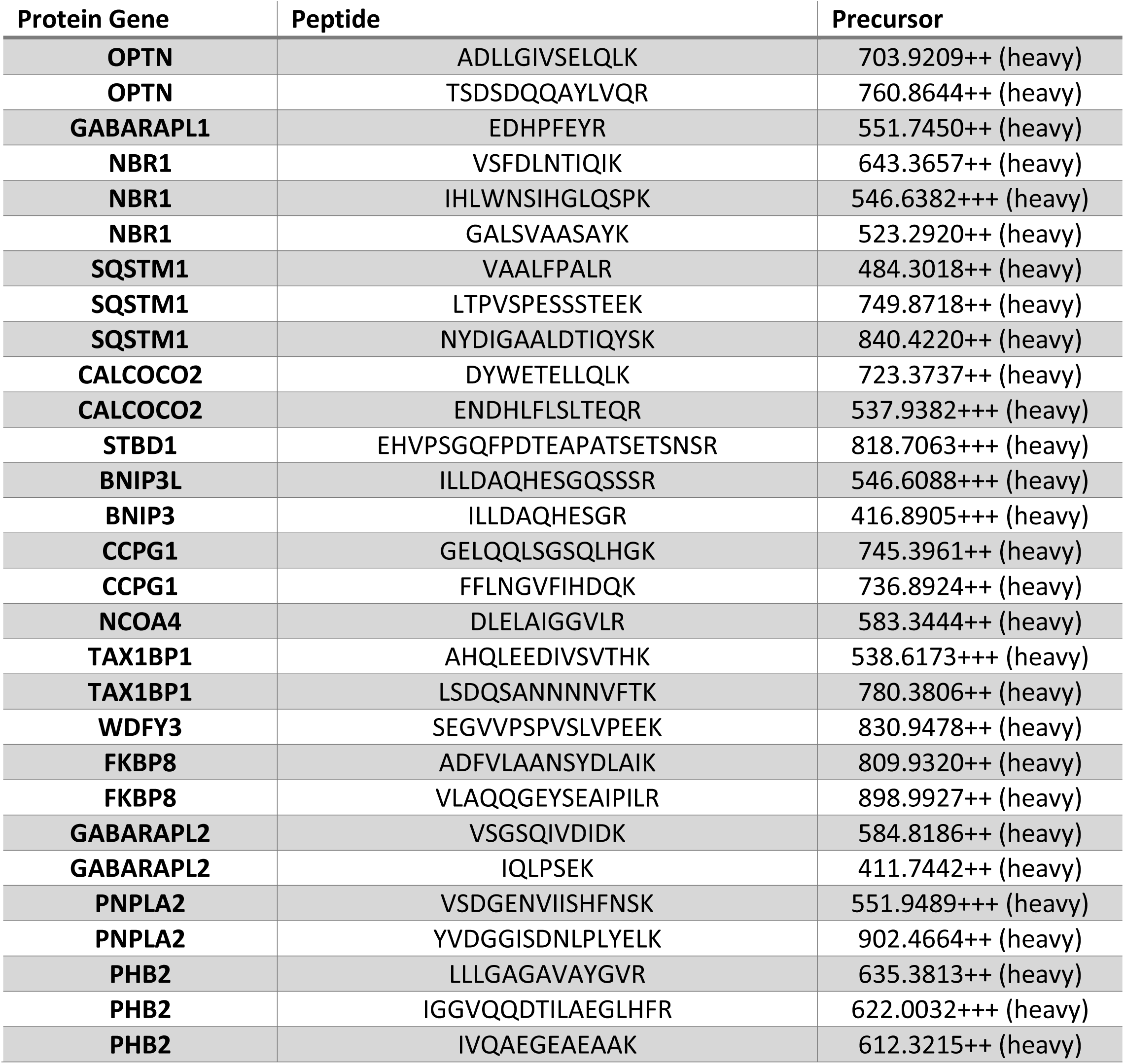
Synthetic heavy-labelled peptides.

#### Targeted Proteomics measurements

LC-MS/MS measurements were performed on a Q-Exactive HF-X mass spectrometer coupled to an EasyLC 1200 nanoflow-HPLC (all Thermo Scientific). 500 ng peptides were separated on a fused silica HPLC-column tip (I.D. 75 μm, New Objective, self-packed with Acquity CSH C18-AQ, 1.7 μm (Waters) to a length of 20 cm) using a gradient of A (0.1% formic acid in water) and B (0.1% formic acid in 80% acetonitrile in water): samples were loaded with 0% B with a flow rate of 600 nL/min; peptides were separated by 4%–30% B within 85 min with a flow rate of 250 nL/min. Spray voltage was set to 2.3 kV and the ion-transfer tube temperature to 250°C; no sheath and auxiliary gas were used. Mass spectrometer was operated in PRM mode with a resolution of 60,000, maximum injection time of 118 ms, AGC target value of 1 x 10^6^ and an isolation window of 1.5 m/z and normalized collision energy of 27. Targeted precursors were acquired in a scheduled manner in 3 min time windows. Loop count was set at 30 and full MS scans were acquired at a resolution of 30,000 with an AGC target value of 1 x 10^6^ and a maximal injection time of 54 ms in a range of 400 to 1200 m/z. The raw MS files were analyzed using Skyline ^123,124^. Precursors were manually filtered for interferences. A minimum of 3 fragment ions were considered for quantification. Quantification values were normalized on heavy peptides and all quantities were extracted from skyline.

### tRNA aminoacylation assay

For the tRNA aminoacylation assay, cells were collected and suspended in a solution containing 0.3 M sodium acetate/acetic acid (NaOAc/HOAc) at pH 4.5. Subsequently, total RNA extraction was carried out using acetate-saturated phenol/CHCl_3_ at pH 4.5 (Thermo Fisher Scientific, AM9720). The isolated RNA was then resuspended in 10 mM NaOAc/HOAc at pH 4.5. The samples were divided into two portions: one-half (2 μg) underwent oxidation with 50 mM NaIO_4_ in 100 mM NaOAc/HOAc at pH 4.5 for 15 min, while the other half (2 μg) was incubated in 50 mM NaCl in 100 mM NaOAc/HOAc at pH 4.5 for 15 min. Glucose (100 mM) was used to quench the reactions for 5 min at RT, followed by purification in G50 columns (Cytiva, 27533001) and precipitation with ethanol. The tRNAs were deacylated in 50 mM Tris-HCl at pH 9 for 30 min at 37°C. After precipitation, the RNA was ligated to the 3′ adaptor tRNA using T4 RNA ligase 2 (NEB, M0351L) for 2 h at 37°C. Reverse transcription was performed with the SuperScript IV synthesis kit (Thermo Fisher Scientific, 18091050). Relative aminoacylation levels were determined by qRT– PCR using tRNA-specific primers.

### Isolation of polysome-associated mRNA

For polysome profiling, cells were resuspended in lysis buffer (20 mM Tris-HCl, pH 7.8, 100 mM KCl, 10 mM MgCl2, 1% Triton X-100, 2 mM DTT, 100 μg/mL cycloheximide, 1X complete protease inhibitor). The cytosolic fraction was obtained through centrifugation at 1300 g for 10 min. This extract was then carefully layered onto a 7% to 47% linear sucrose gradient and subjected to centrifugation in a SW41Ti rotor (Beckman Coulter) at 36,000 rpm for 2 h at 4°C. Nine fractions were collected from the resulting gradients, and RNA isolation from each fraction was isolated using TRI reagent (Zymo, R2050-1-200) according to manufactureŕs protocol as described briefly above. Reverse transcription and qRT-PCR for *AHR* were performed as mentioned above. Using the cycle threshold (CT) values, the percent (%) distribution for the mRNAs across the gradients were calculated using the ΔCT method.

### Ribosome Profiling

For the ribosome profiling, cells were washed with ice-cold PBS supplemented with 100 µg/mL CHX (Sigma Aldrich, C7698) and RP-Lysis buffer (20 mM Tris-HCL pH 7.5 (Thermo Fisher Scientific, 15567-027), 10 mM MgCl_2_ (Sigma-Aldrich, M2393), 100 mM KCl (Sigma-Aldrich, P405), 1% Triton-X 100 (Sigma-Aldrich, T8787), 2 mM DTT (Sigma-Aldrich, D0631), 100 µg/mL CHX, 1x EDTA-free Complete Protease Inhibitor Cocktail (Sigma-Aldrich, 11873580001)) was added. After lysis, all samples were centrifuged at 6400 rpm, 4°C for 5 min. The supernatant was taken and digested with 1 U/µL RNaseI (Thermo Fisher Scientific, AM2295) for 45 min at RT under rotation. Digested lysates were run through 7% - 47% sucrose gradients using a Beckman Coulter ultracentrifuge and SW41 Ti rotor (Beckman Coulter) with 36,000 rpm at 4°C for 2 h. Monosome fractions were obtained and digested with 1% SDS (Sigma-Aldrich, 05030) and 0.113 µg/µL Proteinase K (Roche, 3115828001) for 45 min at 45°C. Resulting footprint RNA was extracted following a standard Phenol-Chloroform extraction (Zymo Research, R2050-1-200) and size-selected using a 10% denaturing PAGE gel.

RP library construction in brief: Footprint RNA was dephosphorylated using 5 U of T4 PNK (New England Biolabs, M0201S). Subsequently, preadenylated UMI-linkers were ligated to the RNA 3’end using 100 U T4 RNA Ligase 2, truncated K227Q (New England Biolabs, M0351L). Residual linker was eliminated by 25 U 5’Deadenylase and 15 U RecJf for 60 min at 30°C. Ribosomal RNA was subtracted using a biotinylated rRNA oligo pool in 1x SSC buffer (3 M NaCl, 300 mM trisodium citrate, pH 7), which were pull down using MyOne Streptavidin C1 DynaBeads (Thermo Fisher Scientific, 65001). Resulting RNA footprints were reverse transcribed using the SuperScript III First-Strand Synthesis System (Thermo Fisher Scientific, 2232161). cDNA was size-selected using a 8% denaturing PAGE gel. cDNA was circularized by using the CircLigaseII Kit (Lucigen, CL9021K). The samples were subjected to PCR to introduce Illumina i7 indexes, followed by size-selection on an 8% non-denaturing PAGE gel. Resulting sample concentrations were measured with Qubit 3.0 (Thermo Fisher Scientific) using Qubit DNA HS kit (New England Biolabs, M0494S). The final RP libraries were single-end sequenced with a NextSeq2000 P2 system (Illumina). All samples were sequenced at the DKFZ Sequencing Open Lab, associated with the DKFZ Genomics & Proteomics Core Facility.

#### RiboSeq data processing

The FASTQ raw data was provided by the DKFZ Genomics & Proteomics Core Facility. In brief, sample adapters were trimmed using cutadapt (v3.4)^125^ and demultiplexed with barcode_splitter from FASTX-toolkit (v0.0.6)^126^. Fragments smaller than 30 nt were dropped. UMIs extraction was performed using umi_tools (v1.1.1)^127^. By BLAST-Like Alignment Tool (BLAT) (v36x2), rRNA reads were filtered and discarded^128^. The rRNA index for RNA18S5, RNA28S5 and RNA5-8S5 was constructed manually from NCBI RefSeq annotation. Remaining reads were aligned with Spliced Transcripts Alignment using STAR (v2.5.3a)^98^ to the GRCh37/hg19 reference with the following call parameters --outSAMtype BAM Unsorted --readFilesCommand zcat --quantMode TranscriptomeSAM GeneCounts --outSAMmapqUnique 0. Genome browser bigwig tracks were obtained using samtools (v1.15.1) and bedtools (v2.24.0). RPF 5’ density was calculated as previously described in Loayza-Puch et al.^129^.

### Statistical analysis

GraphPad Prism (v9.4.1 or v8.4.3) was used for statistical analysis and statistical presentation unless otherwise specified. In case two conditions were compared, a paired or unpaired two-tailed Student’s t-test was performed. If more than two conditions were compared, a one-way ANOVA followed by a Šídák’s multiple comparisons test was applied. Immunoblot time courses with more than two conditions were compared using a two-way ANOVA followed by a Šídák’s multiple comparisons test. For each experiment the number of replicates and the statistical test applied are indicated in the figure legend. Data are presented as mean ± SEM. *p < 0.05, **p < 0.01, ***p < 0.001, n.s.: not significant.

For bioinformatic analysis, unless otherwise stated, all pairwise comparisons were performed using Kruskal-Wallis and Wilcoxon sum rank tests, and all reported p-values were adjusted using the Benjamini-Hochberg procedure. All analyses were run in R, versions 3.3 and 4.2.2, (https://cran.r-project.org/) and Bioconductor version 3.3 and 3.15 (https://bioconductor.org/). All representations were generated using ggplot2, ggpubr, gridExtra and RcolorBrewer.

## Generation of schematic representations

The schematic representations in the graphical abstract were generated using Biorender.org.

## Contact for reagent and resource sharing

Further information and requests for resources and reagents should be directed to and will be fulfilled by the lead contact, Kathrin Thedieck.

## Materials availability

All unique materials and reagents generated as part of this study are available from the lead contact with a completed Material Transfer Agreement.

## Acknowledgements

We thank the patients from the Heidelberg University Hospital and the tissue bank of the National Center for Tumor Diseases (NCT), the department of Neuropathology Heidelberg, especially Ulrike Vogel. We acknowledge Wilhelm Palm and Rafael Paschoal de Campos for scientific discussions and help with the macropinocytosis assays and Stefan Pusch for cloning and plasmids. We acknowledge the DKFZ Core facilities: Light Microscopy Facility (LMF), especially Damir Krunic (for the help with analysis, microscope introduction), the NGS and OpenLab Core facility (RNAseq). We thank Julia Sundheimer, Nicholas Zacharewski, Tim C. Kühn and Alessa L. Henneberg for their help with harvests and immunoblotting. We thank Verena Panitz for critical reading of the manuscript.

KT acknowledges support from the European Union European Research Council (ERC AdG BEYOND STRESS, grant agreement No 101054429) and from the European Partnership for the Assessment of Risks from Chemicals PARC (Grant Agreement No 101057014), and CO acknowledges support from the European Union European Research Council (ERC CoG CancAHR, grant agreement No 101045257), which all have received funding from the European Union’s Horizon Europe research and innovation programme. KT acknowledges support from the PoLiMeR Innovative Training Network (Marie Skłodowska-Curie grant agreement No. 812616) and ARDRE (Marie Skłodowska-Curie grant agreement No. 847681), which have received funding from the European Union’s Horizon 2020 research and innovation programme; the DFG (German Research Foundation (project No TH 1358/3-2), project FG-2000 from the Austrian Science Fund (FWF), and from Stichting TSC Fonds (calls 2015 and 2017). MTP (2019) and KT (2017) are recipients of the Research Award of the German Tuberous Sclerosis Foundation. KT was recipient of a Rosalind-Franklin-Fellowship of the University of Groningen (2013-2019). ST, CO and KT acknowledge support from the BMBF e:Med initiative GlioPATH (01ZX1402). CS and KT acknowledge support from the BMBF e:Med initiative MAPTor-NET (031A426A/B). CS, CO, KT, and BvdE acknowledge support from the MESI-STRAT project (grant agreement No 754688). CH, CO and FS acknowledge support from the German Research Foundation (SFB1389 UNITE-Glioblastoma; project No. 404521405). AS and SM acknowledge support from the German Academic Exchange Service (DAAD). MS was a Research Fellow of the F.R.S.-FNRS (Belgium; Grant No 1.A.385.16). BVdE acknowledges support from the EOS consortium DECODE (Belgium, Grant No 30837538). FL-P acknowledges support from the ERC-STG DualRP project (Grant agreement ID: 759579). HS was supported by grants from the Deutsche Forschungsgemeinschaft (DFG) (INST 337/15-1, INST 337/16-1 & INST 152/837-1). CH was supported by the Deutsche Forschungsgemeinschaft (DFG) (No. 262133997). TK acknowledges Fellowships from Uehara Memorial Foundation and the International Medical Research Foundation. VIK acknowledges grants from BBSRC (BB/M023389/1, BB/R008167/2) and RESETageing H2020 grant (952266). AK was supported by the Austrian Science Fund (FWF) (P 36299) and the Phospholipid Research Center Heidelberg (AKO-2019-070/2-1). AMH acknowledges support by the Tyrolean Science Fund (TWF; grant agreements F.33468/7-2021). JD was supported by the Canton and the University of Fribourg as part of the SKINTEGRITY.CH collaborative research project and by the Swiss National Science Foundation (grant CRSII5_189952). MK acknowledges support by Tyrolean Science Fund (TWF; grant agreement F.18903) and the University of Innsbruck (Project No. 316826).

Views & opinions are those of the authors.

## Author contributions

PH, LH, MTP and PRN designed and performed experiments, analysed data, and contributed to manuscript writing. MS conducted and analysed experimental data. AS designed, performed and wrote bioinformatics methods and analyses for cancer transcriptome and RPPA data. UR designed and conducted kinase assays. AH designed experiments and contributed to scientific discussions. SS and LR designed and performed experiments and analysed data. TB developed MALDI-MS Imaging methods, performed MSI experiments and wrote the corresponding methods. AL and JD conducted targeted proteomics of autophagy regulators. ASE developed, performed and analysed intracellular Trp measurements and analysed proteome data. BB, LFSP, MR, DS, VP, TK, MH, JRP, ILK, TB, SO, LET, FH, AR, DS, IK, CB, MRP and LEE performed and supported experiments and analyses. ST performed analysis of regulatory regions. AvP and YZ supported proteome sample preparation and data analysis. SK measured the Trp stress proteome, supervised by HS. FL-P and AK performed ribosome profiling. LW established and LW and AG performed LC-MS lipid analyses and analysed lipid MS data. TK and VIK performed autophagy gene analysis, and supported autophagy data analysis. JZ supported data analysis. SM performed analysis of MALDI-MS imaging. PS, FS performed tissue collection and pathological assessment. AK supervised lipid measurements by LC-MS. CH supervised MALDI-MS Imaging. MK designed and supervised proteome and intracellular Trp measurement and analysis. CS designed and supervised EGFR and RAS experiments and LAMP stainings. BVdE analysed data and contributed to scientific discussions. CO and KT conceived and supervised the study, designed and analysed experiments, and wrote the manuscript. All the authors read, revised, and approved the manuscript.

